# LncRNA *OIP5-AS1* is overexpressed in undifferentiated oral tumors and integrated analysis identifies as a downstream effector of stemness-associated transcription factors

**DOI:** 10.1101/285270

**Authors:** Ganesan Arunkumar, Shankar Anand, Partha Raksha, Shankar Dhamodharan, Harikrishnan Prasanna Srinivasa Rao, Shanmugam Subbiah, Avaniyapuram Kannan Murugan, Arasambattu Kannan Munirajan

## Abstract

Long non-coding RNAs (lncRNAs) play an important role in the regulation of key cellular processes in early development and in cancer. LncRNA *Oip5-as1* facilitates stem cell self-renewal in mouse by sponging mmu-miR-7 and modulating NANOG level, yet its role in cancer is less understood. We analyzed *OIP5-AS1* expression in oral tumors and in TCGA datasets. We observed overexpression of *OIP5-AS1* in oral tumors (*P*<0.001) and in tumors of epithelial origin from TCGA. *OIP5-AS1* expression was strongly associated with undifferentiated tumors (*P*=0.0038). *In silico* analysis showed miR-7 binding site is conserved in mouse and human *OIP5-AS1*. However, human *NANOG* 3’-UTR lost the binding site for hsa-miR-7a-3. Therefore, we screened for other miRNAs that can be sponged by *OIP5-AS1* and identified six potential miRNAs and their downstream target genes. Expression analysis showed downregulation of miRNAs and upregulation of downstream target genes, particularly in undifferentiated tumors with high-level of *OIP5-AS1* suggesting that *OIP5-AS1* could post-transcriptionally modulate the downstream target genes. Further, systematic epigenomic analysis of *OIP5-AS1* promoter revealed binding motifs for MYC, NANOG and KLF4 suggesting that *OIP5-AS1* could be transactivated by stemness-associated transcription factors in cancer. Overexpression of OIP5-AS1 in undifferentiated oral tumors may confer poor prognosis through maintenance of cancer stemness.

## Introduction

Head and neck squamous cell carcinomas (HNSCCs) include malignant tumors which arise from the squamous epithelial cells of the oral cavity, nasopharyngeal cavity, paranasal sinuses, salivary glands, and larynx. HNSCC constitutes the sixth most common cancer worldwide with an average 5-year survival rate <50% [1]. Oral cancer is the commonest cancer in the South-East Asia and ranks number one among all other cancers in India with approximate rate 32%-40% of total malignancies diagnosed each year in the Indian subcontinent. The chronic use of chewing tobacco, betel quid, areca nut, stalked lime and paan are common in India present in 90% of cases and has been strongly associated with an increased risk of oral cancer [2]. In addition to the tobacco associated carcinogens, several other factors and viruses mainly certain types of Human Papilloma Virus (HPV) may also play a crucial role in oral tumorigenesis [1].

Oral cancer originates as a result of the multi-step process with a multifactorial etiology involving various genetic and molecular changes. Oral epithelial cells are affected by various genetic alterations including mutations, Indels and SNV in *TP53, NOTCH1, PIK3CA, PTEN, RAS, CASP8, RB, FAT1, TRAF3* and *CDKN2A* [3,4,5]. Various genes of the JAK/STAT, RAS/MAPK and PI3K pathways and, apoptotic gene *CASP8* are frequently reported to harbor hot spot mutations in Head and neck cancers [3, 6]. However, mutations in *TP53, PIK3CA, NOTCH1* and *CASP8* were less frequent in South Indian oral tumors [7, 8]. In addition, Epithelial-mesenchymal transition (EMT), a transdifferentiating mechanism that directs changes in cell states along the epithelial versus mesenchymal axes is a major event in oral cancer progression and metastasis. EMT confers epithelial-mesenchymal plasticity upon epithelial cells and the cellular transformation process is orchestrated by a group of transcription factors (TFs), such as the SNAIL, TWIST, and ZEB families [9]. Recently, we reported the role of natural antisense transcript (NAT) for EMT signaling in oral tumors [9]. Therefore, the pathways of carcinogenesis in Indian population might be different from other population which might lead through chromosome instability, telomere lengthening, hormonal activation, environmental factors, chromatin modification, epigenetic changes and dysregulation of non-coding RNAs.

Of late, increasing number of publication showed dysregulation of non-coding RNAs (ncRNAs) in various cancers. Non-coding RNAs are non-protein coding RNA transcripts classified into, microRNAs (miRNAs), a class of shorter ncRNAs with length of ∼17-24 nucleotides involved in the regulation of gene expression at post-transcriptional level and long non-coding RNAs (lncRNAs) which are >200 nucleotides in length with limited or no protein-coding capacity, are involved in various cellular mechanism like RNA/DNA, RNA/RNA and RNA/Protein interaction based gene regulation [10]. Deregulation of miRNAs are often reported in many cancers including oral cancer and some of the miRNAs serve as potential biomarkers for detection of early cancer development, disease reoccurrence and prognosis [11]. However, the role of lncRNAs in oral cancer is less understood and increasing number of novel lncRNA transcripts add on the layer of complexity in understanding their role in cancers.

A few lncRNAs including *MEG3, PTENP-AS1, PANDAR* and *GAS5* functions as tumor suppressors and several other lncRNAs are well established to function as oncogenes in various cancers [12]. *HOTAIR, linc-ROR, H19, CCAT1, ZEB1-AS1, NEAT1, MALAT1, UCA1* and *CDKN2B-AS1* are well documented to be upregulated in oral cancer [12,14,15]. LncRNAs function by acting as signals, decoys, guides, and scaffolds and as a repressor or activator of gene transcription and translation. *cryano* (*linc-oip5*) is a long intergenic ncRNA, first identified in Zebrafish has been reported to be overexpressed in morula stage of the mouse embryo and maintains stem cell self-renewal in embryo through modulation of NANOG by sponging *Nanog* targeting miR-7a [16, 17]. The human counterpart of *cryano* is *OIP5-AS1*, which is transcribed from *OIP5* gene in antisense orientation and its function in human cancer is yet to be explored. Therefore, in this study, we analyzed the expression of *OIP5-AS1* and selected *OIP5-AS1* sponged miRNAs and their downstream target genes in sixty oral tumor tissues. We also carried out *in-silico* epigenomic analysis to understand the function of *OIP5-AS1* in oral tumorigenesis.

## Results

### Patient clinical characteristics

A total of 60 oral squamous cell carcinoma tumors and 8 normal tissue samples were used to study the expression of lncRNAs *OIP5-AS1*. The mean age of the oral cancer patients is 52.75 ±11.12 and males were predominant (71.6%, n=43). Eighty-one percent of the oral cancer patients were tobacco abusers (n=49) and 11 (65%) and 17 female oral cancer patients used tobacco in smokeless form. Only 18% (n=11) of the individuals developed cancer without the habit of oral cancer associated any risk factors. Eighty percent of the patient presented high tumor grade (>T2 stage) and 96% of them were node positive. About 66% (n=44) of the patients had undifferentiated cellular pathology.

### LncRNA OIP5-AS1 is overexpressed in oral tumors with undifferentiated cellular pathology

LncRNA *OIP5-AS1* was found to be overexpressed in oral tumors (*P* <0.001) (Figure 1A) and most of the clinical-pathological characters such as age, tobacco abuse, tumor and nodal stages showed an association with *OIP5-AS1* expression (Figure 1B-D). Only cellular differentiation status showed statistical significance (*P* = 0.0038) with the *OIP5-AS1* expression level (Figure 1E). Fisher exact test did not show any statistical significance, however, undifferentiated oral tumors displayed higher odds ratio (OR) (2.579 [0.8245 – 8.067]) (Table 1). Over 67% of the tumors with undifferentiated pathology expressed *OIP5-AS1* at a higher level. Further, high-grade tumors with the undifferentiated cellular pathology had significant upregulation (*P* = 0.0038) of *OIP5-AS1* than high-grade tumors with differentiated cellular pathology (Figure 1F). Similarly, undifferentiated tumors with tobacco chewing/smoking history showed significant overexpression of *OIP5-AS1* (*P* = 0.0094) (Figure 1G). These results suggest an association of *OIP5-AS1* with cellular differentiation status of oral tumors.

**Figure 1.**
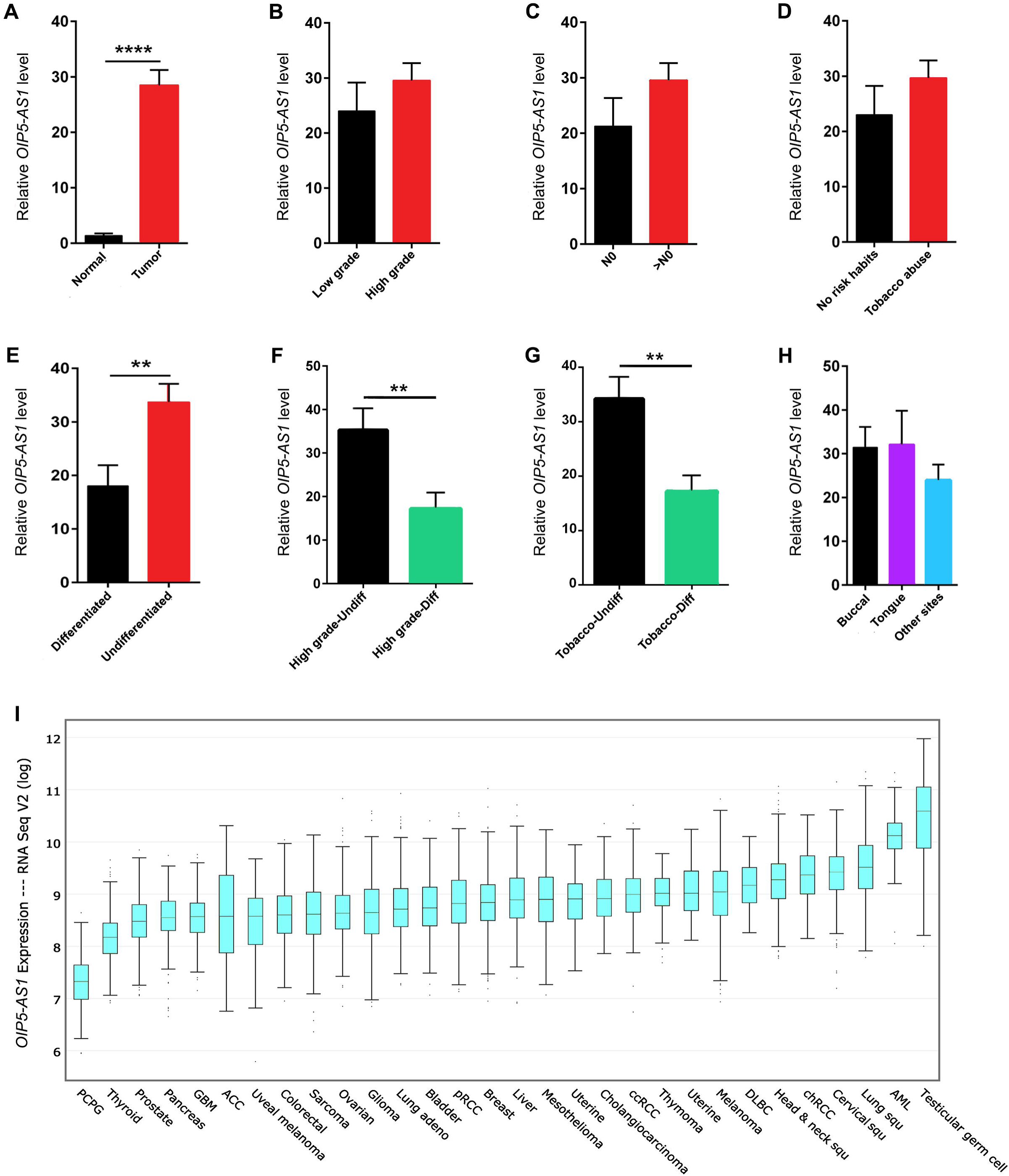
Expression analysis of *OIP5-AS1* in various human cancers. **(A)** Relative expression level of lncRNA *OIP5-AS1* in oral tumors compared with normal tissues. **(B)** Expression level of *OIP5-AS1* between tumor grade, **(C)** tumor nodal stage, **(D)** oral cancer risk habits, **(E)** tumor cell differentiation pathology. **(F)** Relative expression level of *OIP5-AS1* between differentiated and undifferentiated oral tumors in high tumor grade, **(G)** in patients having tobacco abuse. **(H)** Expression of OIP5-AS1 between the tumor sites. Statistical significance represented as ** for *P*<0.01 and **** for *P*<0.0001 (two-tailed Student’s t-test). **(I)** Expression pattern of *OIP5-AS1* in the spectrum of cancers from TCGA database. Testicular germ cell tumors expressed lncRNAs *OIP5-AS1* at high level followed by AML and cancers of squamous epithelial origin including head and neck. Data points are presented in log values.

**Table 1.** Relationship between OIP5-AS1 expression and clinicopathological characteristics in oral cancer patients.

| Clinical characters |  | Number | OIP5-AS1 expression level |  | *P* - value<sup>#</sup> | OR (95% CI) |
| --- | --- | --- | --- | --- | --- | --- |
|  |  |  | Low (n=33) | High (n=27) |  |  |
| Age | <53 | 22 | 12 | 10 | 0.7822 | 1.360 |
|  | ≥54 | 32 | 15 | 17 |  |  |
| Gender | Male | 43 | 25 | 18 | 0.5666 | 1.563 |
|  | Female | 17 | 8 | 9 |  |  |
| Risk habits | Positive | 11 | 7 | 4 | 0.7391 | 1.199 |
|  | Negative | 49 | 26 | 23 |  |  |
| Tumor stage | ≤T2 | 12 | 7 | 5 | 0.9999 | 1.077 |
|  | >T2 | 48 | 26 | 22 |  |  |
| Nodal invasion | Positive | 8 | 5 | 3 | 0.7193 | 1.429 |
|  | Negative | 52 | 28 | 24 |  |  |
| Differentiation | Differentiated | 20 | 14 | 6 | 0.1680 | 2.579 |
|  | Undifferentiated | 40 | 19 | 21 |  |  |

<sup>#</sup> - Fisher's exact test with two tailed *P* value.

### OIP5-AS1 overexpression is common in human cancers of epithelial origin

To check the *OIP5-AS1* expression level in different cancers, we analyzed the human cancer data sets from The Cancer Genome Consortium (TCGA) (Figure 1I). We found a high level of *OIP5-AS1* expression in tumors of epithelial origin, particularly in Lung, cervical and head and neck cancers. Lung squamous cell carcinoma expressed a significantly higher level of *OIP5-AS1* than lung adenocarcinoma. Similarly, renal cell carcinomas also showed overexpression of *OIP5-AS1* suggesting its association with the cancers of epithelial origin. Further, testicular germ cell tumors expressed this lncRNAs at very high level in comparison to all other tumors.

### Human NANOG is not post-transcriptionally regulated by OIP5-AS1

In the mouse embryo, overexpression of *Oip5-as1* (1700020I14Rik) was shown to maintain stemness by regulating the steady state level of NANOG through sponging of mmu-miR-7a [16–18]. We hypothesized that *OIP5-AS1* might have a similar function in human cancers. Therefore, we checked the conservation of hsa-miR-7-5p and mmu-miR-7 mature sequence and observed 100% sequence similarity between mouse and human (Figure 2A). Further, when checked for the conservation of human *OIP5-AS1* sequence with mouse *Oip5-as1*, we observed poor conservation. However, the binding site for miRNA miR-7 was conserved in *OIP5-AS1* (Figure 2B, Supplementary Figure S1).

**Figure 2.**
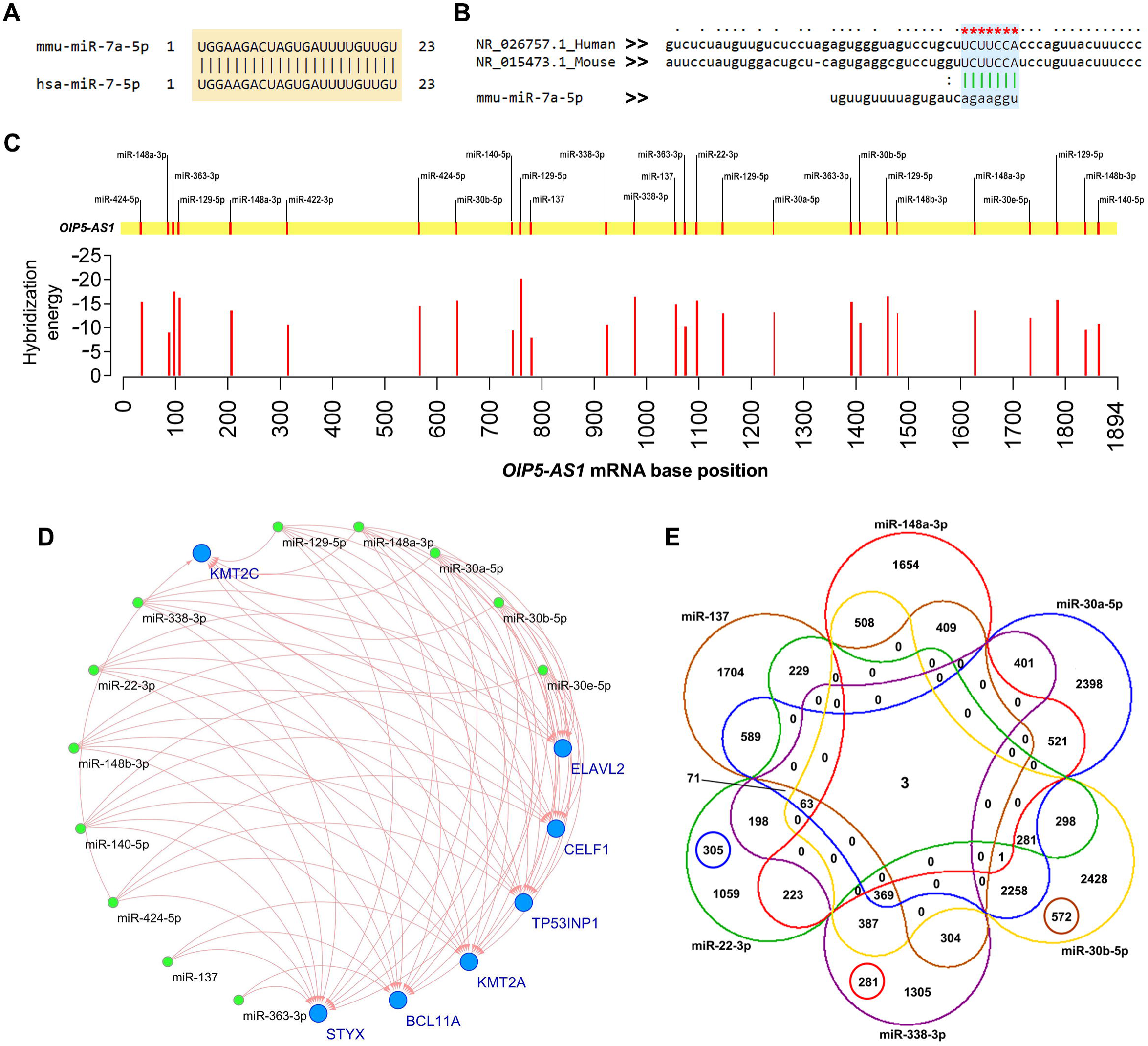
Prediction of miRNA targeted by *OIP5-AS1* and possible interactions. **(A)** Nucleotide alignment of mature sequences of mouse mmu-miR-7a-5p (top) and hsa-miR-7-5p (bottom) shows highly conserved. **(B)** Crustal alignment of human *OIP5-AS1* (top) and mouse *Oip5-as1* (middle) and the interacting site of miR-7 (human/mouse) (bottom). Green lines represent the miRNA seed sequence interaction with the lncRNA and red star indicating the conserved interaction site in human/mouse lncRNA for the candidate miRNA binding. **(C)** Position of predicted miRNA binding sites in the *OIP5-AS1* transcript (top) and their corresponding hybridization energy (kcal/mol) with the base position of the seeds (bottom). The full-length transcript of *OIP5-AS1* (1894 nt) has multiple interaction sites for the miRNAs. **(D)** Interaction network of predicted miRNAs (nodes in green) and their downstream target mRNAs (nodes in blue). **(E)** Six-set Venn diagram of the miRNAs selected in this study and number of shared mRNA targets. *BCL11A, KMT2A* and *STYX* were commonly targeted by all the 6 miRNAs. miR-30a-5p, miR-30b-5p, miR-338-3p and miR-22-3p shared maximum common downstream targets.

In mouse, mmu-miR-7a-5p binds to the 3’-UTR of *Nanog*, regulates its steady state level and signals for cellular differentiation. Overexpressed *Oip5-as1* maintains stemness by sponging the mmu-miR-7a-5p [16]. Since the hsa-miR-7a mature sequence is 100% conserved, we checked whether a similar mechanism is taking place in humans. We did a clustal alignment of mouse *Nanog* and human *NANOG* mRNA and found that human *NANOG* 3’-UTR is not conserved and thus no binding site for hsa-miR-7 (Supplementary Table S1). Moreover, when TCGA HNSCC datasets were analyzed for *OIP5-AS1* and *NANOG* co-expression using ChIPBase v2, we did not observe any significant correlation (*r* = 0.1863).

### OIP5-AS1 can sponge miRNAs with tumor suppressive function

Since human *NANOG* is not regulated by hsa-miR-7a-5p due to lack of binding sites, we asked the question whether the *OIP5-AS1* could maintain the cellular stemness by sponging other miRNAs that can post-transcriptionally control stemness associated TFs factors. We did bioinformatics analysis for miRNAs that can be regulated through sponging by *OIP5-AS1* and their target genes. We collected the *OI5-AS1* target miRNAs predicted by Starbase, DIANA tools and miRcode and identified twelve miRNAs which were targeted by *OIP5-AS1* through interaction sites (Figure 2C).

For experimental validation in oral tumors, we narrowed down that candidate miRNAs to six (miR-137, miR-148a-3p, miR-30a-5p, miR-30b-5p, miR-338-3p and miR-22-3p) by reviewing the functional evidence present in the literature, analyzing their expression in HNSCC datasets from TCGA and correlating with *OIP5-AS1* expression (Supplementary Table S2). For expression studies, along with the 6 miRNAs, miR-140-5p which were shown to control the tumor metastasis in HNSCC and miR-181a-3p, an oral cancer-specific upregulated miRNA that was included as a background control to study lncRNA *OIP5-AS1*’s sponging activity. (*For convenience hereafter, we use “*miR*” to refer human miRNAs*)

### OIP5-AS1 sponged miRNAs targets chromatin modifiers and RNA binding proteins

To ascertain whether the miRNAs targets stemness associated genes, we predicted the downstream target genes for the 12 miRNAs using TargetScan and miRanda online tools. The results showed that seven genes *BCL11A, CELF1, KMT2A, ELAVL2, KMT2C, STYX* and *TP53INP1* were targeted by all miRNAs (Figure 2D, Supplementary Table S3). Further, we constructed a six-set Venn diagram to identify common targets shared by the short-listed 6 miRNAs and identified *BCL11A, KMT2A* and *STYX* as a common target for all the predicted miRNAs (Figure 2E). However, except miR-137 other miRNAs target all the shortlisted genes.

Similar to the miRNA selection, we reviewed the literature for the association of 7 target genes with cancer and analyzed their expression pattern in HNSCC from TCGA database with reference to *OIP5-AS1* expression level (Supplementary Figure S2). *KMT2A, KMT2C* and *BCL11A* were significantly upregulated in the samples that overexpressed *OIP5-AS1* (*P* = 0.0001, 0.0001 and 0.0001, respectively). We also checked the correlation between the 7 target genes with OIP5-AS1 and found correlation rank *r* of 0.4383, 0.4289, 0.456 and 0.3384 respectively with significant *P* – values for *CELF1, KMT2A, KMT2C,* and *TP53INP1* (Supplementary Figure S3). We also tested the co-expression pattern of OIP5-AS1 with the candidate genes in HNSCC cancer cell line, FANTOM 5 and GTEx studies. The *CELF1, KMT2A,* and *KMT2C* showed a positive correlation with OIP5-AS1 expression in many cell types. Further, to understand the functional interactions, we screened for the RNA-binding protein (RBP) binding sites in the 7 target gene transcripts using CLIPdb. The *CELF1, KMT2C,* and *KMT2A* had a maximum number of RBP interactions (763, 485 and 296, respectively) and binding sites (37, 34 and 35 respectively) (Supplementary Table S4). In addition, we analyzed the histone modification (HM) signature at candidate gene loci from Roadmap Epigenomics Project HM ChIP-seq data, gene expression alteration in TFs loss/gain of function from Gene Expression Omnibus (GEO) and ChIP-Enrichment analysis of stemness associated TFs binding at gene promoter. Gene loci of *CELF1, KMT2A,* and *KMT2C* showed maximum HM signature with significant expression alteration during TFs perturbation and enriched stemness associated TFs binding in various cell types along with *OIP5-AS1* (Supplementary Figure S4, Supplementary Table S5). Based on the above results, we selected the three target genes *CELF1, KMT2A,* and *KMT2C* for gene expression studies in oral tumors.

### Downregulation of OIP5-AS1 sponged miRNAs results in upregulation of downstream target genes

The expression of 6 miRNAs and 3 target genes were analyzed in sixty oral cancer samples. Out of the 8 selected miRNAs, miR-137, miR-140-5p, miR-148a-3p, miR-30a-5p and miR-338-3p were significantly downregulated in the tumors compared with normal tissue (*P* <0.001, <0.001, 0.001, 0.001 and 0.0003, respectively) (Figure 3A). We further analyzed the expression of miRNAs with reference to cellular differentiation status and observed low-level expression of miRNAs in undifferentiated tumors, and only miR-22-5p and miR-30b-5p expression were statistically significant (*P* = 0.0485 and 0.0440, respectively). However, the oral cancer specific miRNA, miR-181a-3p was upregulated in the undifferentiated oral tumors (Figure 3B).

**Figure 3.**
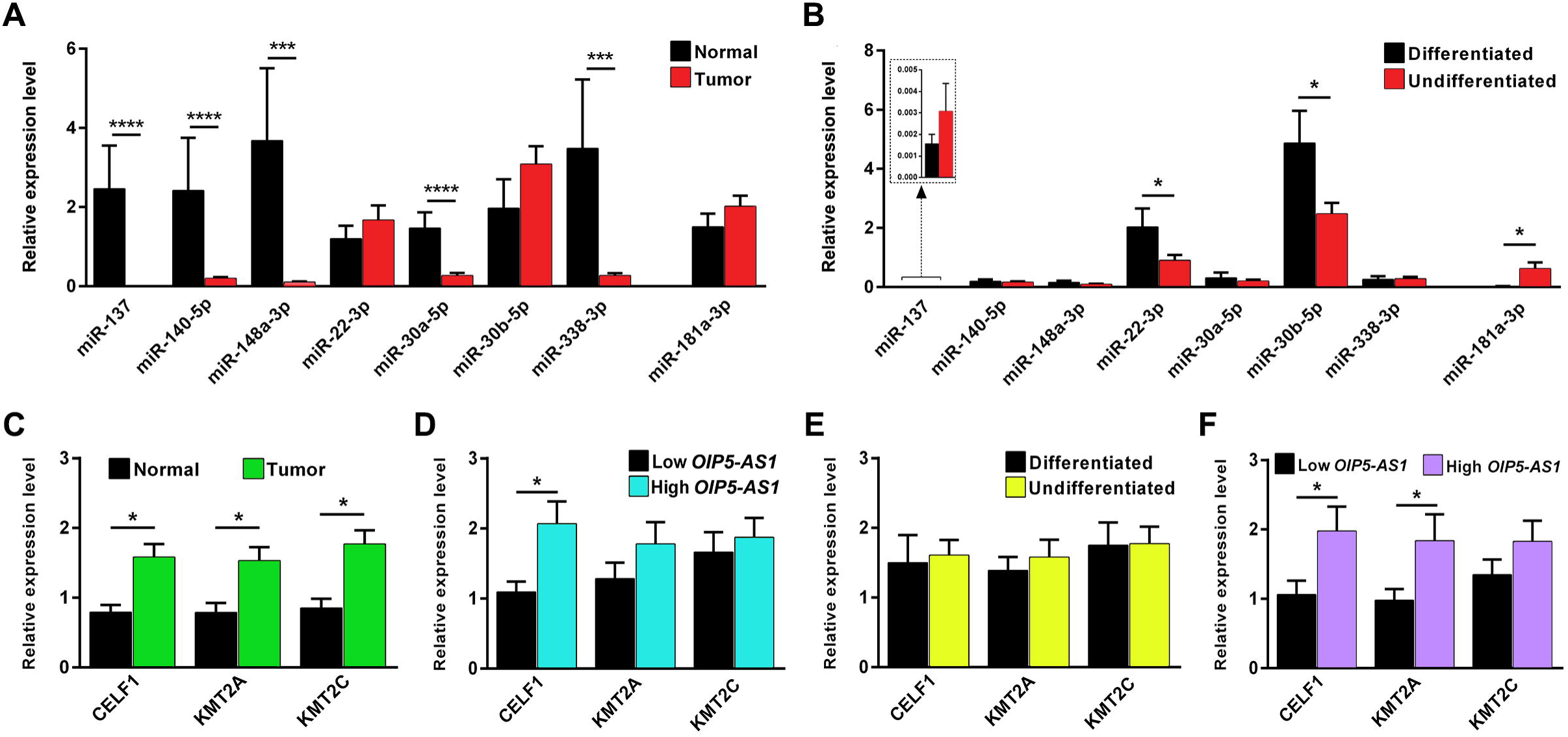
Expression profiling of predicted miRNAs and their downstream target genes. **(A)** Expression levels of predicted miRNAs in oral tumors and normal tissues. Except for miR-22-3p and miR-30b-5p, other miRNAs are significantly downregulated in oral tumors. **(B)** Expression levels of predicted miRNAs in oral tumors between cell differentiation status. miR-137 which shares the least common downstream target and miR-181a-3p which don’t have interaction site with *OIP5-AS1* and nor seed match for stemness TFs was upregulated in undifferentiated oral tumors. **(C)** Expression of *CELF1, KMT2A* and *KMT2C* were significantly overexpressed in oral tumor compared with normal tissues. **(D)** Expression levels of *CELF1, KMT2A* and *KMT2C* in oral tumors that overexpressed *OIP5-AS1* compared to *OIP5-AS1* low expressed tumors. All the 3 mRNAs were upregulated in *OIP5-AS1* overexpressed oral tumors with *CELF1* having statistical significance. **(E)** *CELF1, KMT2A* and *KMT2C* were overexpressed in oral tumors with undifferentiated cellular pathology. **(F)** Undifferentiated oral tumors with high levels of *OIP5-AS1* also upregulated *CELF1, KMT2A* and *KMT2C* compared with undifferentiated tumors with low *OIP5-AS1*. *CELF1* and *KMT2A* which has the maximum number of predicted miRNAs binding overexpressed significantly than the *KMT2C* with the minimum number of predicted miRNAs binding. Statistical significance represented as ** for *P*<0.01, *** for *P*<0.001 and **** for *P*<0.0001 (two-tailed Student’s t-test).

The candidate genes *CELF1, KMT2A* and *KMT2C* were significantly upregulated in oral tumors (*P* = 0.0417, 0.0248 and 0.018, respectively) (Figure 3C). Tumor samples with a high level of *OIP5-AS1* showed overexpression of the above genes than the tumors that expressed low levels of *OIP5-AS1*. Only *CELF1* expression was statistically significant (*P* = 0.0226) (Figure 3D). Further, undifferentiated oral tumors expressed a higher level of candidate genes compared to differentiated tumors (Figure 3E). Similarly, undifferentiated oral tumors with high levels of *OIP5-AS1* overexpressed *CELF1, KMT2A,* and *KMT2C*. *CELF1* and *KMT2A* showed statistically significant expression (*P* = 0.0306 and 0.0341, respectively) (Figure 3F).

### OIP5-AS1 is under the control of Yamanaka factors

Since *OIP5-AS1* is expressed at high level in undifferentiated tumor cells, we asked the question whether the promoter of *OIP5-AS1* gene has possible binding motifs for stemness associated TFs (Yamanaka factors) MYC, KLF4, OCT4, SOX2, and NANOG. Therefore, we analyzed the ChIP sequence data from ENCODE database/Roadmap Epigenomics project for TF bindings in both 1kb up and downstream sequence from transcription start site (TSS) of *OIP5-AS1* gene. We found a strong association of MYC binding with *OIP5-AS1* promoter region (Figure 4A). Surprisingly, we also identified several TF binding motifs for MYC, KLF4 and NANOG in the *OIP5-AS1* promoter through sequence prediction (Figure 4B, Supplementary Table S6). In addition, we also found several binding motifs for MAX, a functional associate for MYC with which MYC forms heterodimer for DNA binding, in the *OIP5-AS1* promoter. Further, to understand the transcriptional activity of the *OIP5-AS1* promoter, we analyzed chromatin modification signatures. We observed strong signatures of H3K4me3 and H3K27ac in hESC and cancer cell lines suggesting that the gene was actively transcribed in stem cells and cancer cells. Further, FAIRE-Seq (Formaldehyde-Assisted Isolation of Regulatory Elements) analysis revealed strong regulatory elements binding around the NANOG motif in hESC cell line and MYC motif in cancer cell lines suggesting that NANOG might regulate *OIP5-AS1* in hESC and MYC in cancer cells. In addition, H3K27ac/H3K9ac active signature in iPS cells also confirms that *OIP5-AS1* could be a downstream effector of Yamanaka factors to maintain stemness (Supplementary Figure S4).

**Figure 4.**
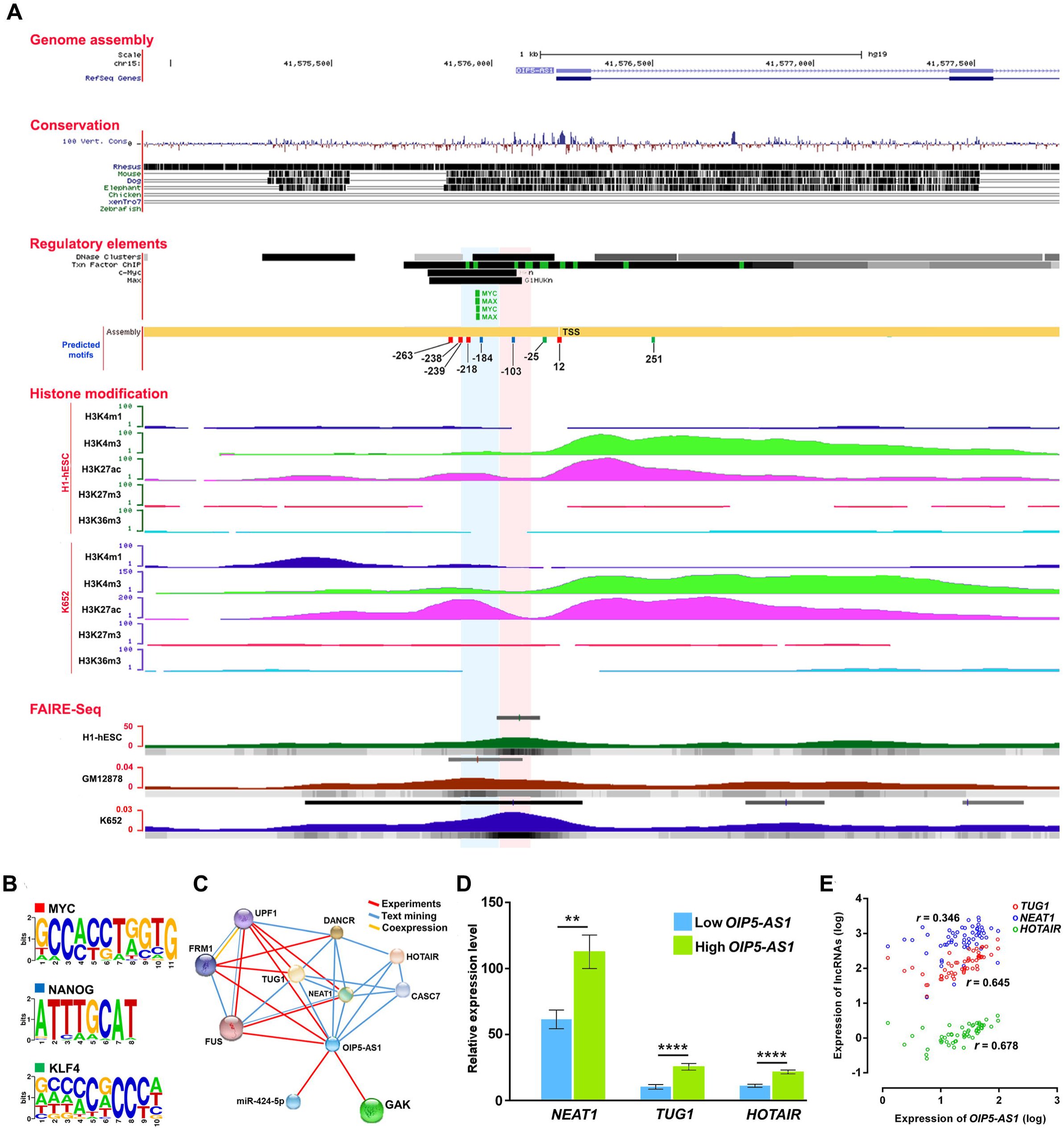
Epigenomic analysis of *OIP5-AS1* promoter and expression of associated lncRNAs. **(A)** Graphical view of *OIP5-AS1* promoter region with regulatory elements binding, histone modification signatures in H1-hESC and immortalized undifferentiated chronic myelogenous leukemia cell line (K562) and position of MYC (red), NANOG (blue) and KLF4 (green) motifs present around 1kb up and downstream from TSS. H3K4me3 and H3K27ac signature are strong in hESC and cancer cells suggesting its active role in maintaining stemness in stem cells and cancer cells. Further, FAIRE-Seq (Formaldehyde-Assisted Isolation of Regulatory Elements) displayed strong TF binding around the NANOG motif in hESC and MYC motif in cancer cells suggesting that the stemness regulatory pathways is facilitated by NANOG in hESC and MYC in cancer cells. **(B)** Sequences of MYC, NANOG and KLF4 motifs screened in the promoter sequence of *OIP5-AS1*. **(C)** Interaction network of *OIP5-AS1* through bioinformatics screening showing *OIP5-AS1* interacts with *NEAT1, TUG1* and *HOTAIR*. **(D)** Expression levels of lncRNAs *NEAT1, TUG1* and *HOTAIR* between *OIP5-AS1* overexpressed and underexpressed oral tumors from this study. Statistical significance represented as ** for *P*<0.01 and **** for *P*<0.0001 (two-tailed Student’s t-test). **(E)** Correlation of expression between *NEAT1, TUG1* and *HOTAIR* with *OIP5-AS1*. *TUG1* and *HOTAIR* have a significant correlation in expression with *OIP5-AS1*.

### OIP5-AS1 could modulate Yamanaka factors through its interacting lncRNAs

As *OIP5-AS1* is not directly regulating genes strongly associated with stemness, we analyzed the *OIP5-AS1* and other lncRNAs/gene interaction network using RAIN online tool and observed the interaction of the *OIP5-AS1* with established chromatin modifiers lncRNAs *HOTAIR, TUG1,* and *NEAT1* (Figure 4C). We checked their expression in oral tumors and in HNSCC from TCGA database. Interestingly, all the 3 lncRNAs were upregulated in the oral tumors that overexpressed *OIP5-AS1* and showed significant positive correlation with *OIP5-AS1* expression (*HOTAIR* – *r* = 0.678, *TUG1* – *r* = 0.645 and *NEAT1* – *r* = 0.346) in TCGA HNSCC datasets (Figure 4D & 4E). There is no report in RAIN database for any functional interaction with Yamanaka factors for the above lncRNAs. However, all the lncRNAs harbor binding sites for miR-143/145 which regulates the Yamanaka factors suggesting that these lncRNAs can collectively maintain stemness in tumors by modulating the Yamanaka factors (Supplementary Table S7).

### OIP-AS1 target genes form a functional network

Finally, to understand the importance of the *OIP5-AS1* regulated gene function at the molecular level, we constructed a protein-protein interaction network using interaction data from BioGRID database. All the targets showed many individual functional interactions and several common interaction partners between KMT2A, KMT2C, CELF1, ELAVL2 and TP53INP1 (Figure 5). The interaction network showed that the target genes are interacting with several RBPs, chromatin regulatory genes, kinase pathway genes and cancer associated genes. ELAVL1 and BMI1 exhibited a maximum shared interaction between the candidate genes. ELAVL1 shares interactions commonly with the selected candidate genes CELF1, KMT2A and KMT2C. The KMT2A and KMT2C shared maximum interacting partners while CELF1 has maximum interactions with other candidate genes through interacting partners.

**Figure 5.**
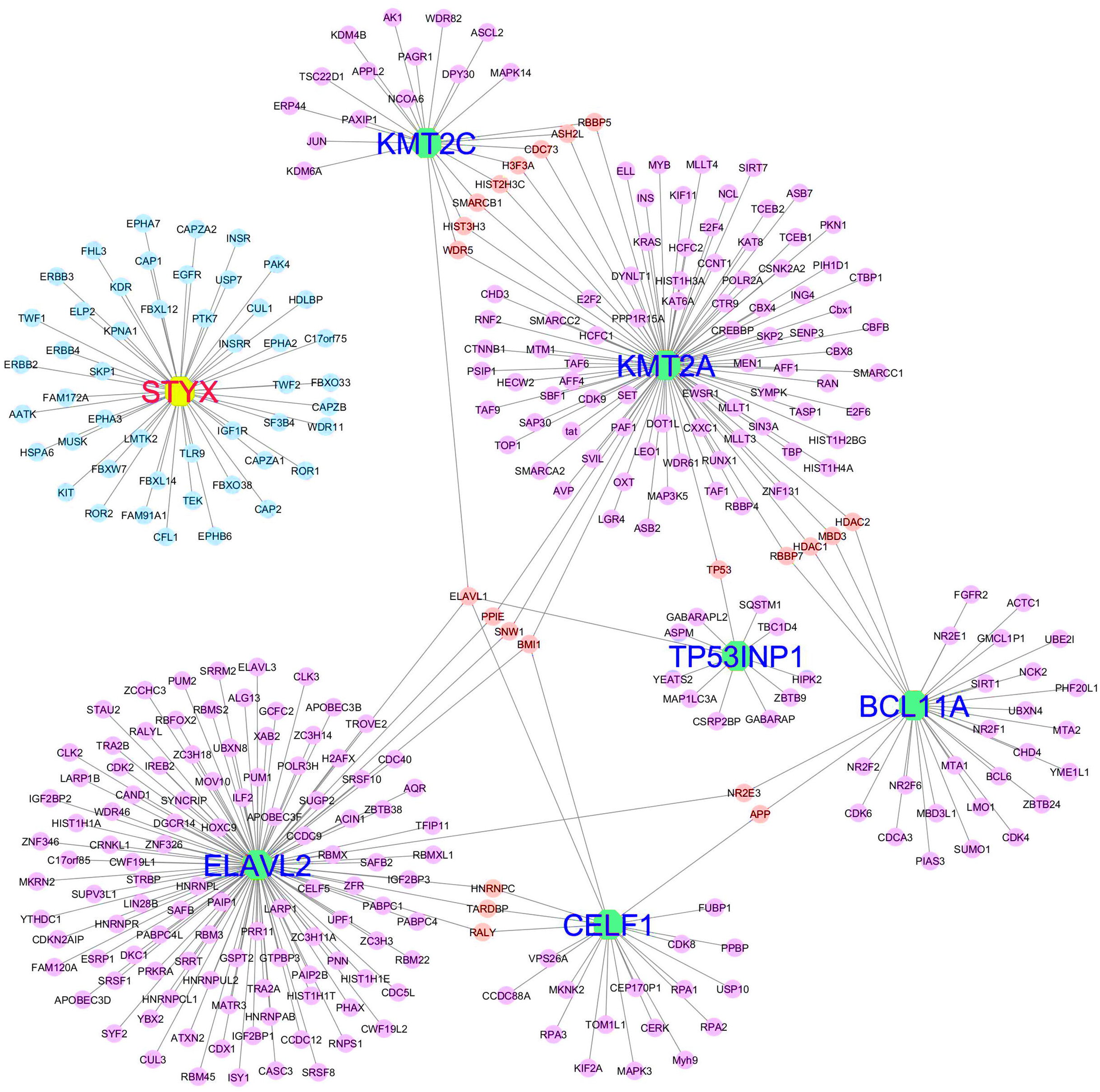
Protein-Protein interaction networks of genes regulated by *OIP5-AS1* sponged miRNAs. The protein interaction network showing the ELAVL2, KMT2A, KMT2C, TP53INP1, CELF1, BCL11A and STYX interaction with various other proteins. Shared interactions are labelled as red nodes and unshared as violet nodes. ELAVL1 and BMI1 were found to be the shared interaction with maximum candidate genes. ELAVL1 shares interactions commonly with the selected candidate genes CELF1, KMT2A and KMT2C. KMT2A and KMT2C shared maximum interacting partners while CELF1 has maximum interactions with other candidate genes through interacting partners. STYX does not share any interaction (blue nodes) with other candidates.

### OIP5-AS1 complements with OIP5 in maintaining stemness

The *OIP5-AS1*, an intergenic lncRNA transcribed in antisense orientation of *OIP5* gene is overexpressed in oral tumors. We checked the expression of *OIP5* in all cancer datasets available in TCGA database. Similar to *OIP5-AS1*, we observed overexpression of *OIP5* gene in squamous cell carcinomas including HNSCC and also in germ-cell tumor (Supplementary Figure S5). In co-expression analysis, *OIP5* was one of the top-most gene found to be co-expressed with *OIP5-AS1* in HNSCC than any other cancer (*r* = 0.5727, *P* < 0.0001) (Supplementary Figure S6). Interestingly, unlike *OIP5-AS1*, *OIP5* is not having motifs with stemness TFs binding to its promoter and the activation was independent. Further, we screened the 3’-UTR of *OIP5* for any shared microRNA response element (MREs) for the miRNAs those having interaction with *OIP5-AS1* and found only miR-424-5p and miR-30a-5p specific MREs in *OIP5*. Interestingly, we also found many MREs for stemness regulatory miRNAs miR-143/145 family, EMT regulatory miRNA miR-200a/miR-200b/miR-141, TP53 induced miR-34a, let-7, and several other oral cancer specific tumor suppressive miRNAs in *OIP5* mRNA 3’-UTR (Supplementary Table S7). These results suggest that *OIP5* co-expression could complement *OIP5-AS1* function in maintaining stemness in oral cancer.

## Discussion

With the advent of next generation sequencing, an abundance of non-coding RNAs have been discovered outnumbering the protein coding RNA transcripts. Non-coding RNA transcript is divided into two major class; microRNAs and long non-coding RNAs depending on the length of the mature transcripts. miRNAs are small non-coding RNAs exert their function at the post-transcriptional level by binding to the 3’-UTRs of the coding mRNAs via a unique seed sequence. Aberration in miRNAs expression was associated with various cancers and diseases has been documented [11,19,20]. Deregulated miRNA expression could alter the post transcriptional regulation of the target genes. One such important function of miR-200 family is regulation of epithelial to mesenchymal transition and miR-143/145 is to maintain the stemness [9, 14]. EMT is an important process by which epithelial cells acquire mesenchymal, fibroblast-like properties during development and accumulating evidence points to a critical role of EMT-like events during tumor progression and malignant transformation by giving the cancer cells the metastatic potential [21]. Synergistically, Cancer stem cells (CSCs), a small niche of cells with self-renewal capacity within the tumor can play an important role in field cancerization [22]. The role of miRNAs in the regulation of EMT and stemness in development and cancer is indispensable; however, the lncRNA’s role in cancer development is yet to be explored.

LncRNAs, with less or no protein coding potential, are shown to be involved in early embryonic development to cancer metastasis [12, 23]. *OIP5-AS1*, a long intergenic ncRNA transcribed in antisense from the *OIP5* gene, located at chromosome 15q15.1, was first discovered as *cyrano* in zebrafish and reported to function as a major regulator of neurogenesis during development and in the maintenance of self-renewal in mouse morula stage modulating the *nanog* expression by sponging mmu-miR-7 [16, 17]. The function of this lncRNA in human development/cancer is not clear and there are only a couple of studies highlighting its role in human cancers. Therefore, we analyzed the expression of *OIP5-AS1* in oral cancers of South Indian origin.

In this study, we found overexpression of *OIP5-AS1* in oral tumors with undifferentiated cellular pathology and in tumors from the patients with tobacco chewing/smoking history. Our results are consistent with HNSCC datasets from TCGA database and notably, cancers from the squamous epithelial origin and testicular germ cell expressed very high levels of *OIP5-AS1*. A similar high expression level of *OIP5-AS1* has been reported in glioma and shown to be associated with advanced tumor stage and promoted tumor migration through down-regulation of YAP-NOTCH signaling pathway [24].

In mouse, overexpressed *Oip5-as1* sponge the cell differentiation promoting mmu-miR-7a-5p and maintains stemness modulating the *nanog* level. Therefore, we checked the binding sites for the mouse miRNA counterpart in human (hsa-miR-7-5p), as this particular miRNA is highly conserved across the vertebrates [17]. Though human *OIP5-AS1* was poorly conserved, the binding site for hsa-miR-7-5p is preserved at a conserved part of the transcript. To the surprise, the 3’-UTR of human *NANOG* did not carry any binding site for hsa-miR-7-5p suggesting that *NANOG* was not targeted by hsa-miR-7-5p. Therefore, we performed a bioinformatics screening and identified 12 miRNAs interacting with *OIP5-AS1*. Six miRNAs miR-137, miR-148a-3p, miR-338-3p, miR-30a/b-5p and miR-22-3p known to be associated with several cancers were chosen to study the expression levels in oral tumors [20,25,26]. Interestingly, all these miRNAs were also reported to be dysregulated in neuronal development and psychiatric disorders [19, 27].

Expression profiling of the 6 shortlisted miRNAs revealed that most of the miRNAs were downregulated in oral tumors and miR-22-3p and miR-30b-5p were significantly downregulated in undifferentiated tumors. Our results suggest that the predicted miRNAs which are established as tumor suppressors were downregulated in oral tumors and this could be due to the sponging effect of *OIP5-AS1*. Further, in undifferentiated tumors, *OIP5-AS1* alone or together with other lncRNAs might sponge miR-22-3p and miR-30b-5p to a greater extent resulting in the derepression of the downstream target genes. miR-140-5p which has been shown to inhibits cell migration in hypopharyngeal carcinoma did not show any change with reference to tumor differentiation status [28]. Moreover, the commonly upregulated oral cancer specific miR-181a-3p which does not have interaction site with *OIP5-AS1* remains upregulated (*P*=0.0319) in undifferentiated tumors suggesting that the other miRNAs might be tightly regulated by overexpressed *OIP5-AS1* through its sponging activity.

Further, we screened the downstream target genes of the 6 miRNAs to check whether any of them targets *NANOG*. None of the miRNAs could target *NANOG* or any other well-known stemness related Yamanaka factors (*OCT4, SOX2, KLF4,* and *MYC*), however, were found to target epigenetic, metabolic, transport and other development related genes [29]. Seven genes, *CELF1, KMT2A, KMT2C, ELAVL2, STYX, BCL11A,* and *TP53INP1* were the common gene targets for all the 6 miRNAs. We narrowed down the candidates to three (*CELF1, KMT2A,* and *KMT2C*) using a six-set Venn diagram of miRNAs with additional parameters like the number of target sites for the 6 miRNAs in 3’-UTR, binding score, the number of RBP interacting sites and reviewing the literature.

We found overexpression of the candidates *CELF1, KMT2A,* and *KMT2C* in tumors and particularly in undifferentiated oral tumors. Further, tumors with high levels of *OIP5-AS1* also overexpressed the candidate genes and specifically in undifferentiated tumors. CELF1, a RNA-binding protein, binds to GU-rich element (GRE), a regulatory sequence in mRNAs, controls a grid of mRNA transcripts that regulate cell division, proliferation, and apoptosis. Transcripts coding for suppression of cell growth and proliferation are actively targeted by CELF1 protein and upon binding to them, designated for degradation in malignant T-cell [30]. RNA-sequencing identified 1283 mRNA transcripts involved in various cellular pathologies and are differentially regulated due to overexpression of CELF1 and promoted alternative splicing in oral cancer cells [31]. CELF1 protein was reported to be significantly overexpressed in human breast cancer tissues by functioning as a central node controlling translational activation of genes driving EMT and tumor progression [32]. Overexpression of CELF1 was reported to prevent apoptosis by destabilizing pro-apoptotic mRNAs in oral cancer cells [33]. Loss of *CELF1* expression downregulated the MAPK signaling pathway and promoted colorectal cancer cell proliferation and chemoresistance [34]. *KMT2A* and *KMT2C* are lysine methyltransferase protein coding genes of mixed-lineage leukemia (*MLL*) family which encode the nuclear protein with an AT hook DNA-binding domain zinc finger also reported as one of the highly mutated genes in genome level. Dysregulation or mutation of the *KMT2* family changes the epigenetic identity of the cells and drives a subset of infantile and adult leukemia [35]. KMT2A contain the CxxC domains that bind to nonmethylated CpG dinucleotides that are highly enriched around TSS and required for active non-coding transcription at enhancers. KMT2A is preferentially expressed in glioma stem cells and downregulation reduces CSC self-renewal and tumorigenicity [36]. KMT2A has been found to interact with the NF-κB pathway to regulate brain cancer growth and promotes melanoma growth by activating the hTERT signaling [37]. Knockdown of KMT2A suppressed tumorsphere formation and the expression of cancer stem cell markers [37]. Similarly, KMT2C is also shown to play roles in metastasis of esophageal squamous cell carcinoma and knockdown experiments showed EMT-like morphological change in pancreatic cancer cell lines [38]. Decreased expression of KMT2C was associated with attenuated cell proliferation in pancreatic ductal adenocarcinoma and poor outcome in breast cancer [39, 40]. Enrichment of H3K4me3 at promoters of stemness genes OCT4, Nanog and Sox2 were reported during erythroblast erythroid differentiation but completely lost at upon erythroid differentiation [41]. H3K27me3 and H3K4me3 double-positive signals are involved in cell stemness [42]. Therefore, the overexpression of KMT2A and KMT2C could maintain the chromatin active signature and facilitate the maintenance of stemness in cancer stem cells. In addition, overexpressed CELF1 may account for alternative splicing of pre-mRNAs resulting in dominant negative isoforms and destabilization of pro-apoptotic mRNAs there by giving efficiently to the cancer stem cells to create resistance for chemo/radiotherapy.

Since *OIP5-AS1* was overexpressed in undifferentiated tumors and associated with maintenance of self-renewal in stem cells, we analyzed the *OIP5-AS1* gene promoter region to understand its transcriptional control. We first confirmed the active transcription of the gene by analyzing histone modification signature and DNase-I signature which showed a hyperchromatin activity and hypersensitivity, respectively in hESC and cancer cell lines suggesting the active transcription of *OIP5-AS1* during development and cancer. We found several motifs for MYC, NANOG and KLF4 binding in the upstream of TSS and ENCODE data confirmed the binding of TFs, particularly MYC to the *OIP5-AS1* promoter. Besides, we also found several motifs for MAX, a known myc-associated factor X protein that forms a heterodimer with MYC and transcriptionally active MAX/MYC heterodimer promotes cell proliferation [43]. In addition, we found strong regulatory binding elements near MYC motif in *OIP5-AS1* promoter in undifferentiated myelogenous leukemia cell line. Moreover, we previously reported the upregulation of Yamanaka factors in oral tumors with undifferentiated pathology and *MYC* co-expression with overexpression of lncRNA *CCAT1* in patients with a poor therapeutic response [14, 15]. These findings suggest that the overexpression of *OIP5-AS1* might be due to the transcriptional activation of *OIP5-AS1* by the stemness associated transcription factors.

In addition, we also screened the expression of established chromatin modifying lncRNAs *NEAT1, HOTAIR,* and *TUG1* [44], and observed a significant upregulation in oral tumors that expressed a high level of *OIP5-AS1*. Further, all 3 lncRNAs harbored interaction sites for miRNAs targeting Yamanaka factors, especially for miR-143/145 suggesting that overexpressed *NEAT1, HOTAIR* and *TUG1* could modulate the post-transcriptional control of the stemness factors by sponging miR-143/145. These transcription factors could in turn bind to the *OIP5-AS1* promoter and transactivate the gene resulting in overexpression of *OIP5-AS1*. Overexpressed *OIP5-AS1* may sponge the miRNAs and derepress the target genes *CELF1, KMT2A,* and *KMT2C* (Figure 6). Being important players in chromatin modification, deregulated expression of *KMT2A,* and *KMT2C* genes may lead to open chromatin configuration resulting in active transcription leading to malignant transformation. Further, *NEAT1* was reported to bind active chromatin sites and was significantly accounted for changes in transcriptional activity by binding to TSS [45]. *HOTAIR* was reported to acts as a scaffold by providing binding surfaces for several chromatin-modifying complexes and to facilitate H3K27/H3K4 methylation and demethylation signatures [46]. Moreover, *TUG1* was reported to promote self-renewal of glioma stem cells by sponging miR-145 in the cytoplasm and employing polycomb to repress differentiation genes by locus-specific methylation of histone H3K27 in the nucleus [47]. Therefore, the co-expression of lncRNA *NEAT1*, *HOTAIR* and *TUG1* has pivotal role not only in sponging stemness TFs targeting miR-143/145 but also interacts with active euchromatins to maintain stemness properties.

**Figure 6.**
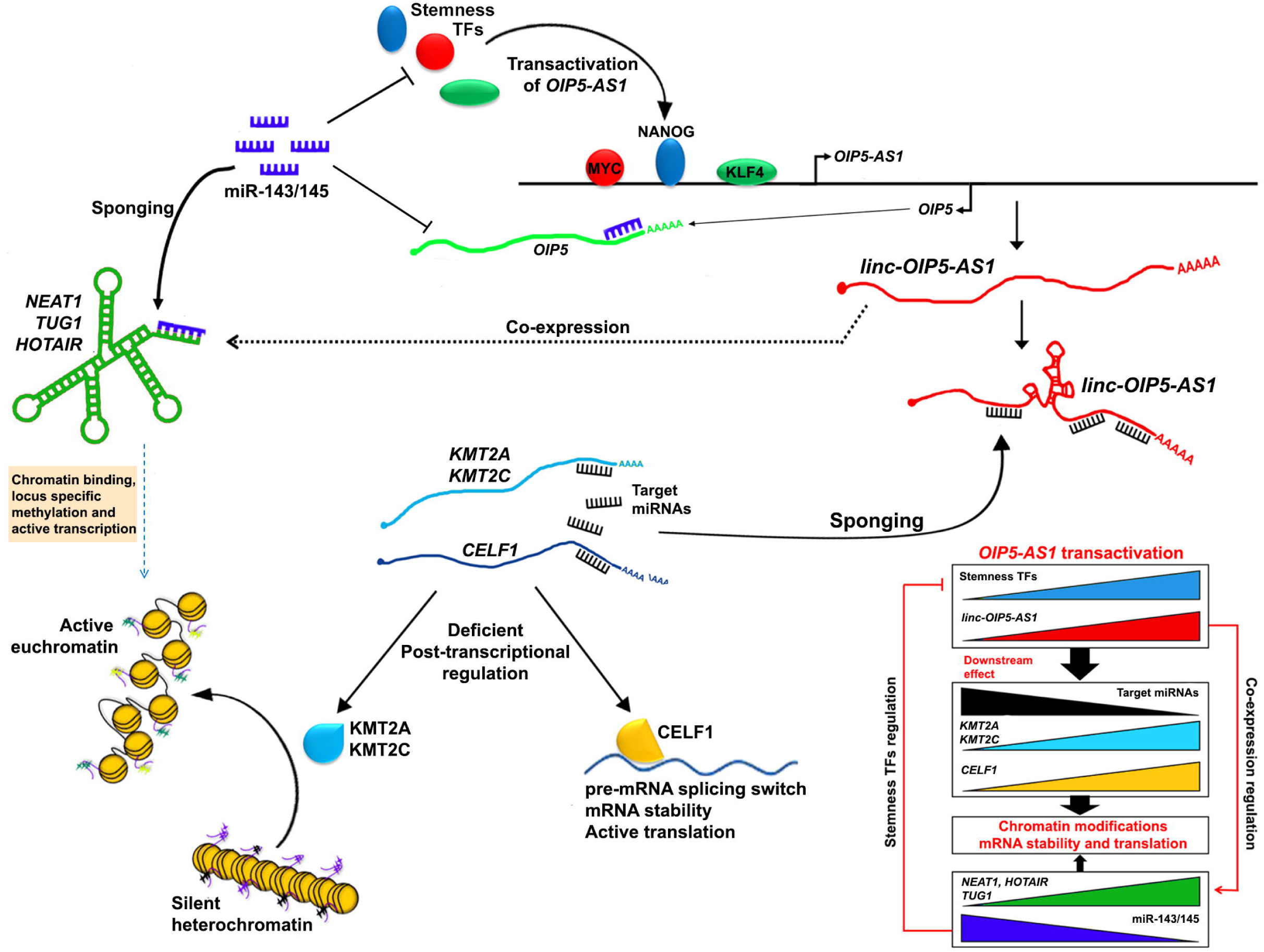
Transactivation of *OIP5-AS1* by stemness regulatory TFs and downstream molecular cascade. Stemness associated TFs transactivates *OIP5-AS1* by binds to the promoter. The overexpressed *OIP5-AS1* sponges the miRNAs targeting *KMT2A, KMT2C* and *CELF1*. With deficient in post-transcriptional regulation, KMT2A, KMT2C and CELF1 overexpress resulting in chromatin modification and increased mRNA stability and active translation specific mRNAs. OIP5-AS1 in addition activates co-express of *NEAT1, TUG1* and *HOTAIR* which sponges stemness regulatory miR-143/145 and maintains the steady state level of stemness TFs. Further, in *cis NEAT1, TUG1* and *HOTAIR* can bind to open chromatins facilitating the binding of KMT2A and KMT2C along with other chromatin modifying enzymes to modify methylation signature and enables active transcription of the genes in the euchromatin sites. The sponging of miR-143/145 also facilitates the overexpression of *OIP5* mRNA.

In addition, *OIP5*, the gene harboring *OIP5-AS1* gene in antisense orientation, was shown to be overexpressed and significantly associated with several cancers [48]. We, therefore, analyzed the *OIP5* gene expression in TCGA HNSCC and observed a significant co-expression with *OIP5-AS1*. *OIP5* did not share miRNA or TFs that regulates *OIP5-AS1* expression level. However, *OIP5* was targeted by stemness regulatory miR-143/145, EMT associated miR-200 family and oral cancer-specific tumor suppressor let-7 family. In oral cancer, the expression of *OIP5* may not be regulated at the post-transcription level as the miR-143/145, miR-200 and let-7 family microRNAs were reported to be downregulated in oral cancer [9,11,14]. These results suggest that with *OIP5* gene, lncRNA *OIP5-AS1* may play a synergistic role in oral tumorigenesis. In summary, our work presented here showed the significant association of *OIP5-AS1* with dedifferentiation pathology in oral tumors. The bioinformatics analysis of lncRNA, miRNA, and mRNA interactions followed by the expression profile identified RNA networks regulating two key chromatin modifiers and one gene involved in the regulation of transcription in oral cancer. Moreover, systematic epigenomic analysis of *OIP5-AS1* suggested that the positive transcriptional regulation of *OIP5-AS1* by co-expressed *NEAT1, TUG1* and *HOTAIR* by recurring miR-143/145 targeting the stemness associated transcription factors might account for the maintenance of stemness and dedifferentiation in tumors and accounts for poor prognosis. Further functional dissection to ascertain the role of *OIP5-AS1* in cellular level is warranted.

## Methods

### Clinical specimens

The present study was approved by the Institutional Ethics Committee (IEC) Madras Medical College, Chennai (No.04092010) and Government Arignar Anna Memorial Cancer Hospital, Kancheepuram (No.101041/e1/2009-2) and was conducted within the ethical framework of Dr. ALM PG Institute of Basic Medical Sciences, Chennai. Sixty oral squamous carcinoma tissue samples and 8 normal tissues were collected from Government Royapettah Hospital, Chennai and Arignar Anna Memorial Cancer Hospital and Research Institute, Kancheepuram. The patient’s contextual and clinicopathological characteristics were documented with standard questionnaire following the IEC guidelines and written informed consent was obtained from each patient, after explaining about the research study. The tumor specimens were collected in RNAlater solution (Ambion, USA) and transported to the laboratory in cold-storage.

### RNA isolation and quality control

Extraction of RNA was carried out as described previously [14]. In brief, tissues were washed twice with ice cold PBS to make it free from residual RNA later solution and homogenized using MicroSmash MS-100 automated homogenizer (Tomy, Japan) with Zirconium beads. Total RNA was isolated using the RNAeasy mini kit (Qiagen, Germany) as per the supplier protocol. The RNA was quantified using NanoDrop2000 UV-Vis spectrophotometer (Thermo Scientific, USA) and the integrity of RNA was verified by resolving in 1% agarose gel in Mupid gel electrophoresis (TaKaRa, Japan).

### cDNA synthesis and quantitative Real-Time PCR

cDNA synthesis was carried out using total RNA (2 μg for mRNA and lncRNAs and 10 ng for miRNAs) and real-time gene expression analysis was performed in ABI Quantstudio 6K Flex (ABI Life Technology, USA) as described in [14]. The list of gene specific primers and miRNA stem loop primers were presented in Supplementary Table S8-S10. *GAPDH* served as an endogenous control for lncRNA and coding genes, and *RNU44* as an endogenous control for miRNAs. Each assay was done in triplicate and the expression level was calculated using 2^−ΔΔCt^ calculation.

### Bio-mining for target prediction, gene interactions, and network construction

For gene expression comparison and identification of target genes from the list of predicted genes cancer datasets from TCGA database was analyzed and downloaded using cBioportal and ChIPBase v2. online tools [49]. Predictions of miRNA targets were performed and retrieved from TargetScan, miRanda, miRcode, Starbase and DIANA online prediction tools. For prediction of RNA-RNA interactions and RNA-RNA binding protein (RBPs) interaction, IntaRNA [50], and CLIPdb [51] online tools were employed. Individual study for assessment of the association of miRNAs in various human cancers was collected from public databases PubMed using MeSH terms such as miRNA/lncRNAs/Gene names, cancer, stemness, EMT, metastasis, prognosis, biomarker, and development. Data from functionally validated studies only were considered for selection of miRNAs and genes. Further, we devised a range of cut-off by reviewing the literature. Hybridization scores for lncRNA-miRNA interaction were calculated using IntaRNA tool. Expression levels and correlation ranking with *OIP5-AS1* expression were calculated based on TCGA HNSCC datasets to narrow down the predicted target miRNAs. Chromatin immunoprecipitation (ChIP) data of human embryonic stem cells and cancer cell lines were accessed through ENCODE, Roadmap and ChIPBase v2 database and UCSC genome browser was used to visualize the gene location and transcript graphic maps. RAIN and BioGRID v3.4 databases were used to screen lncRNAs-RNA/Protein interaction network and validated data of protein-protein interaction respectively [52, 53]. Cytoscape v3 was used for visualizing molecular interaction networks.

### Statistical analysis

The relationship between clinicopathological characteristic features with expression was examined by Fisher exact test and odds ratio (OR) with 95% confidence interval (CI) was calculated to check the risk association. Differences between the means were presented as means ± SEM and analyzed using Student’s *t*-test (Mann-Whitney) using Graph Pad Prism statistical software, v 6.01 (Graph Pad software Inc, USA). Person correlation test was performed for co-expression analysis and *r* rank above 0.3 was considered as a significant association. All tests were two-tailed and a *P* value of <0.05 was considered as statistically significant.

## Acknowledgements

We thank the patients for valuable clinical samples. GA gratefully acknowledges University Grant Commission (UGC), Government of India for providing research fellowships. SA and PR thank Indian Academy of Science for supporting Internship program and providing fellowships. We also gratefully acknowledge the Government of India’s Department of Science and Technology - Fund for Improvement of S&T Infrastructure (FIST), UGC - Special Assistance Programme (SAP) and Department of Health Research – Multi-disciplinary Research Unit (MRU) infrastructural facility.

## Authors’ contributions

AKM (corresponding author) conceived the study designed the experiments and supervised the work. GA contributed to the study design and performed the experiments, analyzed the data and wrote the manuscript. SA and PR carried out bioinformatics analysis. SD assisted the real-time PCR experiments. PSR and SS provided tumor samples and clinical data. AKM analyzed the data and critically reviewed the manuscript. All authors read and approved the final manuscript.

## Ethics approval and consent to participate

The present study was approved by the Institutional Ethics Committee (IEC) Madras Medical College, Chennai (No.04092010) and Government Arignar Anna Memorial Cancer Hospital, Kancheepuram (No.101041/e1/2009-2) and was conducted within the ethical framework of Dr. ALM PG Institute of Basic Medical Sciences, Chennai. The patient’s contextual and clinicopathological characteristics were documented with standard questionnaire following the IEC guidelines and written informed consent was obtained from each patient, after explaining about the research study.

