## Supplementary Materials for "LncRNA *OIP5-AS1* is overexpressed in undifferentiated oral tumors and integrated analysis identifies as a downstream effector of stemness-associated transcription factors"

**Figure S1: Clustal alignment of lncRNA *OIP-AS1*.** Clustal alignment of Zebra fish *cyrano*, Human *OIP5-AS1* and Mouse *Oip5-as1* transcript

```
NR_046387.1_Zebra      cagtagcagcacacgggaccgaaatggcgtaacgcgcagtcgagcaccgcagcagcgcag
NR_026757.1_Human      ---------------------------------------ttg-----agaagct------
NR_015473.1_Mouse      ctttcgcgc-ctccgcccctgggttggcgga-------cagg-----ggaagga------
                                                                *      * **

NR_046387.1_Zebra      aggggccaggctcagcgaacaccacagagcctgg-----cccgaacaacttcaaccagca
NR_026757.1_Human      --------------gcgaagatggcggagtaaggcgtgccgctgcaaactg---------
NR_015473.1_Mouse      --------------gcgaagatggcggagtgaggcgtgccccgggcagctg---------
                                     ***** *   * ***   **     * *    * **

NR_046387.1_Zebra      aatttgttgaaaattactcttcactccagtccacatgtttccaaaccagcacgaga----
NR_026757.1_Human      ---------------------gcctctgggccgggggc-----gagcagcccccgggagg
NR_015473.1_Mouse      ---------------------gcctgcgggcccggggc-----aggcctcccgtgagaag
                                              **   * **    *         *  * *  *

NR_046387.1_Zebra      ----tcaccgctgacgtgccc-ctcatttgtcatatcaagaagagaaaagaacagtcctg
NR_026757.1_Human      ccgagtgcatctgttggaccgtgcgaggagaa---------------------------g
NR_015473.1_Mouse      cggagtacttgtgctggaccgtgtgaggaaccatatcac-------------------ac
                              *   **  *  **     *

NR_046387.1_Zebra      aagggaacacacgaccagatgagaccatcttgtctttgttcttcaaaccgcccttcatcc
NR_026757.1_Human      aaaaaaaatatcggccagaggagaggaact--------------aaccgaacattct---
NR_015473.1_Mouse      aagatgaatattttcctgaccttaggtgctt-----ttatattcagctcaacattct---
                       **    *       ** **    *    **              *      * ***

NR_046387.1_Zebra      tgataagttgaccttcctccagcgttatttcagggaagtgtttgttcag---cataccaa
NR_026757.1_Human      --------------tcctccctaccttatagaggggagtggtcatctacactaaagcaaa
NR_015473.1_Mouse      --------------gaactcaggacttg-gcacgtaagtagatagagataatcatcccaa
                                          *     *     * *  ***        *     *  * **

NR_046387.1_Zebra      gccaagtgttactggtctatgaacatatcatctctgcactttggagaagacaa-ctgaag
NR_026757.1_Human      gt-----gttaggcgcgt---------------c----cgtgcgaagagacca-cca---
NR_015473.1_Mouse      gtcctgttgtagcggctt--gaaagg-----ctc----cat-agaagcgacatacta---
                       *        **   *  *               *    * *  **   ***   *

NR_046387.1_Zebra      ttttttttaattcacataaactcccccacaaactcttcttgtgttcaaaaaagctaaatg
NR_026757.1_Human      ------------------------------aacaggctttgtgttccttatcac---agg
NR_015473.1_Mouse      ------------------------------aact-gt--taagagcaacagaag---atg
                                                     ***      *  *  *   *       * *

NR_046387.1_Zebra      acaacacaattgtgccacaacatggtgccatcgactggtcatattgttcagaataaatca
NR_026757.1_Human      aagaaaatttt----cct------tgaccttta-ggtgcttttatattcatctcaaaaac
NR_015473.1_Mouse      aagataaaaatgagtacaagccagctaccatgg-ctcacctttatgcccagctc------
                       *  * *    *                ** *          *  *   **

NR_046387.1_Zebra      aatatggttgcagtttttgttaaatgaagataaaaatgcagaaaaaaatgcacatttaaa
NR_026757.1_Human      aaaat------------tctgaactcaggac-------ttggcaagt-------------
NR_015473.1_Mouse      ----t------------tgggacctggggaa-------ctga--gga-------------
                           *            *   *  *   **          *

NR_046387.1_Zebra      cagattaa-gtgcctagacgagtctctttttaatagctcctgcctcaatttccatggctt
NR_026757.1_Human      gtttttgaaatgtgg----ctatcacttctctacaca--------ttacctccatgattt
NR_015473.1_Mouse      cag--tgagatgcag----atgtc---tttcgacaga--------ttacctccatggttt
                            * *  **          **   * *  * *            *  ******  **

NR_046387.1_Zebra      cgctctaaagtaaagcacttctttgga--cattttagtctttcaaatggctgaaaatgta
NR_026757.1_Human      gga----atggaaaaggccacttttctttttgttctgcctctcaaattcaacacagagga
NR_015473.1_Mouse      agg----atgcaga-gactgcctttcttttac--------gtcagactcaacacaggaaa
                        *     * * * *   *  * **                 *** *      * *    *

NR_046387.1_Zebra      gtttctgagacgacaaactgtagacagctagctgttctcttaattcacaactcatattgg
NR_026757.1_Human      gct---------------cctaggattccagttatcctgctaac----------------
NR_015473.1_Mouse      gcc---------------gtgaggat-gcagccgtcctgttgac----------------
                       *                    **      **   * **  * *
```

```
NR_046387.1_Zebra      tgctttagcaaaac--catgacccatgctttcatgttttctacagtat--atacttgtac
NR_026757.1_Human      ---atctccaggaagaaagaagcaactcacatgggtcttttgctgtttgcttaattata-
NR_015473.1_Mouse      ---ttcagcaagaa--aacaaaccatggacttgggtcttctgcagctt-ttgaattgga-
                           *   **  *    *  * * *         ** ** * * *  *    * **  *

NR_046387.1_Zebra      tgacaaaccaagacaggcagtggcataacaacccaaaaaaaacataagctttataaatcc
NR_026757.1_Human      ---------aagacatcattttgca-----agcag---aaggctgag-tttca-------
NR_015473.1_Mouse      ---------aagacggtg-tttgca-----agtag---aaggaagatgctgaa-------
                                *****      * ***     *       **     *   *  *

NR_046387.1_Zebra      cacagttaaacctttctagcggggtgctattgagttgcaggatgttatgaaaaatagtta
NR_026757.1_Human      ----tttgaaa---------caggtgctt---aggtggtggtatttgtgaatacttttca
NR_015473.1_Mouse      ----tttgaaa---------caagtgctt---aggttctgtggc--gtgagttgttttct
                            ** **             *****    ** *   *       ***    *  *

NR_046387.1_Zebra      ttttactacctc------------------------------------------------
NR_026757.1_Human      ---------ttccaagcaagaagactaaagaagtagcaagtatgaatgacttcagggttt
NR_015473.1_Mouse      ttttaaaagatccaaacaagaagactgaacaaataccaagtgtgggtgacttcaggtttt
                                 **

NR_046387.1_Zebra      ----------------------aga---------agttaccacagctcagag--ccacac
NR_026757.1_Human      aaaaaaaatgtcttccagtttcagccactaccatgataagcacagttgagactgcagcag
NR_015473.1_Mouse      ttttttaatgtcttccaattttaac----actattataagcacagttga-----ccgcag
                                             *             * * ***** * *     *  **

NR_046387.1_Zebra      ttggatcaaactataagagcatgcttgtctttttttgtttgcagtaaaataaatatagtcg
NR_026757.1_Human      taaattccaaata------------tgtgtttctaatttgacgtgaaagatactaaaaat
NR_015473.1_Mouse      taagttccaaata------------tttaa-tataatttgaagtgaaagaaat-------
                       *    ** ** **            * *   * *  ****  ** *** * *

NR_046387.1_Zebra      ttatag---tctgtctaatggcttggtaatctgatgagaagaggaattaatttttctttt
NR_026757.1_Human      ttatatttgtatatttaaatcctggctcatcctgtgaca---------------------
NR_015473.1_Mouse      ---------gttatctaaattctggctcactctttgaca---------------------
                                  * * ***   ** * * *     *** *

NR_046387.1_Zebra      gtatgtgtgaaatatcgtattgagactgtatgttctccagaataagataaacatcatg--
NR_026757.1_Human      -----------------------------tagatttactgaataggaacaaaggcccaat
NR_015473.1_Mouse      -------------------------------gttttcctggacaaaaacaagatacga--
                                                      * * * * * * *  *  **

NR_046387.1_Zebra      ttcatccattaacgttatgcct--------------------------------------
NR_026757.1_Human      tttaaacaaaaacc-taggccgggtgtggtggctc--acacctgtaatcccaacactttg
NR_015473.1_Mouse      taaaagcagaaa-------ccaggtttgattttgtatgggagtgaaatcattacagtttt
                       *  *  **  **       **

NR_046387.1_Zebra      --gaagaaaggcagtaaaatggttaaatgatcataacatctttaaatagttgctaatagt
NR_026757.1_Human      ggaggccaaggcaggcaaatca---cctgaggttgggagtttgagac---------cagc
NR_015473.1_Mouse      atgggta--------taaatct-------------acatttttaaac---------aaga
                                       ****                 *  ** * *           **

NR_046387.1_Zebra      ctaat-atgttggacagtaagactggtttctagttatctgctatagagcactgtgaa---
NR_026757.1_Human      ctgaccaacatgg---agaaaccccatctctacta----aaaatagaaaattagccagg-
NR_015473.1_Mouse      ttttt-ctcttggagaagaaacctgttgtttataa----gaag----gctttggacagtt
                        *        ***     **  *   * * **                   *    *

NR_046387.1_Zebra      -aataggaatttctcttcatctgt--tgtgacact---------------acatcttttt
NR_026757.1_Human      ---------------------tgtggtggtagata------cgtgtaatcccagcta--c
NR_015473.1_Mouse      taatagtcttaaatgttctgctttgctgttctagtccttctcaggaaggcccagctg--g
                                            * *  **    *                  ** **

NR_046387.1_Zebra      tgttgttgttggtatagt--agttcctatcatacggatggtagatattcaagtgttctga
NR_026757.1_Human      tcgtgaggctgaggcaggagaattgcttgaatccgggaggcggaggttgcagtgaactga
NR_015473.1_Mouse      gcctgtggctagttcag---aat-----------g-------------ggaggaaacata
                          **  * *     **   * *           *               **    *  *

NR_046387.1_Zebra      gttttcatgagaggaagagtagcgaggcctcagtgtgggaaatg-----tgcaatatttg
NR_026757.1_Human      gattgcaccactgcactccagcctgggcgacagagtgagactccgtctcagaaaaat---
NR_015473.1_Mouse      ggctgggacaatgctcaccctgaactgccagcgtgtgggaggccaaggcaggaagagcag
                       *  *     *  *             **    * *** **          * ** *
```

```
NR_046387.1_Zebra       tt-tctttgtcttgttttgagatgaatatgtatattgtacaaacaagtgacaagttgttc
NR_026757.1_Human       ------------------------------------------------------------
NR_015473.1_Mouse       atgttcagggccagcttggacatcaagagg------ctcaaaaccagt-----------a

NR_046387.1_Zebra       gcaaaaacaacaaaat-caccaatgtcttccattaaatgatgtatagtttcatctgcttg
NR_026757.1_Human       ------------------------------------------------------------
NR_015473.1_Mouse       ggaaaaaaaacagagcccagtaacaccctcttttatattcct-tttgtggaatcatgtag

NR_046387.1_Zebra       ctttagggtggcgggatttgtagtgccagcgtagacatggacatgcatgtacggtagcga
NR_026757.1_Human       ------------------------------------------------------------
NR_015473.1_Mouse       cat--gtgagactgggttggaag--------------ggaactggcatatgtggtatt--

NR_046387.1_Zebra       tgcttgctgagtttaatagtcgt------------------agtgacattcagtataggc
NR_026757.1_Human       --------------------------------------------aaaaaacaact-----
NR_015473.1_Mouse       -------tcagttaaggtgatgtggtggaccctgatgtgtaactggccttcacct-----
                                                                          **

NR_046387.1_Zebra       tataataaagtcagttagtacataaa----tatag-gcaaagatttagttagtaaacacc
NR_026757.1_Human       ---------------ttggacatagaggagttgggggcagaggg--gg-gag----gatg
NR_015473.1_Mouse       ---------------ttggacatacaggaatgaaggaaaaa-------------------
                                       * * ***** *    *   *   * *

NR_046387.1_Zebra       ataga-tgcagtatccacaccttggttaatttgattaagaagcgagtagt---agggctg
NR_026757.1_Human       agagaattttccatgtaactccttttccttaagaaaaagaagcaaaaatccctcatgcag
NR_015473.1_Mouse       ctaaagcttggcatataaatcttttcttctaagaaaaagaagcaaagctgtctaatgtag
                          * *       **  *   * *      *  **  ******* *           *  *

NR_046387.1_Zebra       cacgattctggctaaaatg-aaaatcacgtttttttgttgttgttttttttttaatcaag
NR_026757.1_Human       tgcca-tctgactttatggt-gtttcacatta---tatagttactttt--tttaataaac
NR_015473.1_Mouse       cattg-tctgacttttcaacatttttacttta---tgtggttgcattt--ttttataaaa
                              **** **           * ** **    * * ***   ***  *** ** **

NR_046387.1_Zebra       at-cgcgattttctcacaattctatagatgtagaataaaggtttatatgagaaggctatt
NR_026757.1_Human       gcataagattaactcttcctgcaacccaaggtggatacttggatgtt-----------tg
NR_015473.1_Mouse       gtataaa--ta--agttccttcaacccaaaaaaaaaaa----------------------
                                 *         * * *   *      * *

NR_046387.1_Zebra       attattattgttaattttcaaatatatggtaaaacaacattagcattaggcctgtgtcat
NR_026757.1_Human       a---tttttatttatttatttatttattt---------------tttgaggcagtctctc
NR_015473.1_Mouse       ------------------------------------------------------------

NR_046387.1_Zebra       tctaacgttttgtataggcttggaatggtggactggttcatgtctttaacaacattatta
NR_026757.1_Human       tct-----gttgcccaggc---------tggagtgatgcgtgatttctgc----tcactg
NR_015473.1_Mouse       ------------------------------------------------------------

NR_046387.1_Zebra       gaaaatttttttcactggtaaaattaaaccagtagtatcgtcattaatggtctatctaca
NR_026757.1_Human       cagtctctcccacccgggttc-----aagcgat--tctcctgcctca--g--cctcccaa
NR_015473.1_Mouse       ------------------------------------------------------------

NR_046387.1_Zebra       atagataatgttacatgtggaaaattaactaaaatgctcacattcacatgtttgtgcaat
NR_026757.1_Human       gtagctaggattacaggcgcgc-----accaccacgctcagcctgattt--ctag-----
NR_015473.1_Mouse       ------------------------------------------------------------

NR_046387.1_Zebra       aatagaccttttgatttgaatagttttgtggagccctttgtgttgaattcatgcctttta
NR_026757.1_Human       --------------gctgaaaggct------------------gggactcaggtactt--
NR_015473.1_Mouse       ------------------------------------------------------------

NR_046387.1_Zebra       tggtaacctgttgcatgttttctgacgctttagttcgagtgcatatctgaatcagtt---
NR_026757.1_Human       ---tccccagttgggactgaactttctcatcag--agaatgggcctcagaagcaatgttc
NR_015473.1_Mouse       ------------------------------------------------------------
```

```
NR_046387.1_Zebra      ---catttttgttcctgtcaatataatttagtagctataacaaccacaacgatctcttta
NR_026757.1_Human      ccaaacttgtggttgggtcatc--------gtgg-aaaagaaacctcaaata------ag
NR_015473.1_Mouse      ------------------------------------------------------------

NR_046387.1_Zebra      aaagaaaataacaatagtttaaaaaatgagcaacaggaaacattaaaacaattttatata
NR_026757.1_Human      aaaaataattacaat-----aagaaatggattttcttttttcccacaacagtt----ata
NR_015473.1_Mouse      --aaaaa------------------a----------------------------------
                         * * *                  *

NR_046387.1_Zebra      catcaaatag-ccaaattttgctggaaaaatgctcttttggtaacatatttcattcggta
NR_026757.1_Human      cattataaagaacagactgtcgtagaaaactgt-ct-ttgcttccaaatcagcagagg-a
NR_015473.1_Mouse      ------------------------------------------------------------

NR_046387.1_Zebra      taatttgtatctttattttaaagtatttttgttggtcagttttctacaaaataaacgaaa
NR_026757.1_Human      c-cattgtat--gtatt-----------------gtcaggtctttatataa---------
NR_015473.1_Mouse      ------------------------------------------------------------

NR_046387.1_Zebra      agaacgaatgcattttagctgaagtgatgtgcaaatacttcttcaataataagggaatta
NR_026757.1_Human      -gagtgaaac--------------------------------------------------
NR_015473.1_Mouse      ------------------------------------------------------------

NR_046387.1_Zebra      tgaattaaattcaagtaaacctattataacataaaatatagtatttctaataattttatt
NR_026757.1_Human      ------------------------------------------------------------
NR_015473.1_Mouse      ------------------------------------------------------------

NR_046387.1_Zebra      tatttaatagatttgagtttgaattacagtatcacaatactatttgattccacaatactt
NR_026757.1_Human      ------------------------------------------------------------
NR_015473.1_Mouse      ------------------------------------------------------------

NR_046387.1_Zebra      caactggtatagtatcttaacgtaaattaatagtatcgcgacaaccctaataaatatagt
NR_026757.1_Human      ---------------------------------------------cttt-----------
NR_015473.1_Mouse      ------------------------------------------------------------

NR_046387.1_Zebra      atgggacacgtaacagcatttggaggtgtccacattgctgagcaacttgcctaaaacaat
NR_026757.1_Human      ------------------------------------------------------------
NR_015473.1_Mouse      ------------------------------------------------------------

NR_046387.1_Zebra      gtgaagacttgtgcgactgcccttacgcttctctcctgacttaatatacgtaggtgaatc
NR_026757.1_Human      --------------------atttatgcttctt--gtgagtagaga--------------
NR_015473.1_Mouse      ------------------------------------------------------------

NR_046387.1_Zebra      gtacaacagctggattaagatcatgtgtagggtcgaatcaagatcacgttcttttatcga
NR_026757.1_Human      ------------------------------------------------------------
NR_015473.1_Mouse      ------------------------------------------------------------

NR_046387.1_Zebra      ttaatcgtgcagccctagcaagtaagcatttactcttggttttagtttttaagtgacata
NR_026757.1_Human      ------------------------------------------------------------
NR_015473.1_Mouse      ------------------------------------------------------------

NR_046387.1_Zebra      cgctgtagtttattcactggtaatcactattagttgatgataacgtcatagcatgctgag
NR_026757.1_Human      ------------------------------------------------------------
NR_015473.1_Mouse      ------------------------------------------------------------

NR_046387.1_Zebra      gattagtaaaatggcggttggtattgatgtgcgactagccatgctgaaatgcatatatat
NR_026757.1_Human      ------------------------------------------------------------
NR_015473.1_Mouse      ------------------------------------------------------------
```

```
NR_046387.1_Zebra      gtttcataagcactgtgtcagtttgtcctagtttgtactgtgcaaaatgaagttctgggc
NR_026757.1_Human      ------------------------------------------------------------
NR_015473.1_Mouse      ------------------------------------------------------------

NR_046387.1_Zebra      tcaaaatagtcgactcagatcaaaacgtctgtcccaaataggccaaaatttggctcagta
NR_026757.1_Human      ------------------------------------------------------------
NR_015473.1_Mouse      ------------------------------------------------------------

NR_046387.1_Zebra      gcttagaatacgcaggtgttgaagaacttcattcagtacctgtttcaaggttgctaaaac
NR_026757.1_Human      ------------------------------------------------------------
NR_015473.1_Mouse      ------------------------------------------------------------

NR_046387.1_Zebra      tgtactggacagaaagaatagacttctctcttcctaaatagaactgtgtttgttttcata
NR_026757.1_Human      ------------------------------------------------------------
NR_015473.1_Mouse      ------------------------------------------------------------

NR_046387.1_Zebra      cctctaatattgcattctccatcaggtaatgttgtctcaaactagagctgtgggtagatt
NR_026757.1_Human      ------------------------------------------------------------
NR_015473.1_Mouse      ------------------------------------------------------------

NR_046387.1_Zebra      attcctagtagcagaacatgtagattttaactcaacgtgtagaaaattaagtggcataaa
NR_026757.1_Human      ------------------------------------------------------------
NR_015473.1_Mouse      ------------------------------------------------------------

NR_046387.1_Zebra      gcaattatcaaagcattgtaatgtagataaatacctcaatgactggaatgcaaactatat
NR_026757.1_Human      ------------------------------------------------------------
NR_015473.1_Mouse      ------------------------------------------------------------

NR_046387.1_Zebra      atttttgaatatactcgaaaaaatatgcaagccatgatgtgtattaagttcatcttttag
NR_026757.1_Human      ------------------------------------------------------------
NR_015473.1_Mouse      ------------------------------------------------------------

NR_046387.1_Zebra      aagcaataatcagtttatgcatgagttaaagtaccggaacagtttagtggtgtttgtaat
NR_026757.1_Human      ------------------------------------------------------------
NR_015473.1_Mouse      ------------------------------------------------------------

NR_046387.1_Zebra      atctgtagcatatctgtgacagtgacaagtgtttgatcctgaatatctcattgctcataa
NR_026757.1_Human      ------------------------------------------------------------
NR_015473.1_Mouse      ------------------------------------------------------------

NR_046387.1_Zebra      ttaaaagttttccattcctccatttttcatggacaccaaataaatatgttaaggtagcac
NR_026757.1_Human      ------------------------------------------------------------
NR_015473.1_Mouse      ------------------------------------------------------------

NR_046387.1_Zebra      tgggggtctgactaataaagaggattttttgggtgggttgacgaaagttagaagaatcat
NR_026757.1_Human      ------------------------------------------------------------
NR_015473.1_Mouse      ------------------------------------------------------------

NR_046387.1_Zebra      ctacatatgcagggcttttgctcttattaaatttgcatgtttacctaatcattgtggaaa
NR_026757.1_Human      ------------------------------------------------------------
NR_015473.1_Mouse      ------------------------------------------------------------

NR_046387.1_Zebra      gcacataagctgtatgtttattgacagcttattatcattaacagcaatatgagcagtgtt
NR_026757.1_Human      ------------------------------------------------------------
NR_015473.1_Mouse      ------------------------------------------------------------
```

```
NR_046387.1_Zebra      ctgggtagcaatttgtaatactgatgactcattatcctcatgaagcctggacaaacattt
NR_026757.1_Human      ------------------------------------------------------------
NR_015473.1_Mouse      ------------------------------------------------------------

NR_046387.1_Zebra      taccactttacatttacaaaccccatttggagtaaagtatttttttaaagagtgcaaaaaa
NR_026757.1_Human      ------------------------------------------------------------
NR_015473.1_Mouse      ------------------------------------------------------------

NR_046387.1_Zebra      aacctgatgcatggtagaaatggataagaaatgttttgtgctgcatgcaataattgagct
NR_026757.1_Human      ------------------------------------------------------------
NR_015473.1_Mouse      ------------------------------------------------------------

NR_046387.1_Zebra      ttgcttgaaaatgtttatgcagaaatgtgatgacactaaatctatgagtggaccagtgtt
NR_026757.1_Human      ------------------------------------------------------------
NR_015473.1_Mouse      ------------------------------------------------------------

NR_046387.1_Zebra      gtgactttctttttaaatgcatggaatcccaaagttgatctctactgtaaattattcaca
NR_026757.1_Human      ------------------------------------------------------------
NR_015473.1_Mouse      ------------------------------------------------------------

NR_046387.1_Zebra      gacgtgttcattgagaaatgtttgattactgtatgatggaattgtaacaagtttgttctt
NR_026757.1_Human      ------------------------------------------------------------
NR_015473.1_Mouse      ------------------------------------------------------------

NR_046387.1_Zebra      caagaggcaaagtcagatgcagtgttttagcaaagtcttaacaattgtgagttgaaacgg
NR_026757.1_Human      ------------------------------------------------------------
NR_015473.1_Mouse      ------------------------------------------------------------

NR_046387.1_Zebra      gaaaaactatttgtgttatgttcaatgcaaataatattgcagaatataagttaatatgct
NR_026757.1_Human      ------------------------------------------------------------
NR_015473.1_Mouse      ------------------------------------------------------------

NR_046387.1_Zebra      ttgcagacctgatttgcgttgctgattttgcctctgaatgtgaaactctatgcttagtct
NR_026757.1_Human      ------------------------------------------------------------
NR_015473.1_Mouse      ------------------------------------------------------------

NR_046387.1_Zebra      gatctatgcactacagaagcgtcccgcatggttttgtacaaaatattaaattttagcaat
NR_026757.1_Human      ------------------------------------------------------------
NR_015473.1_Mouse      ------------------------------------------------------------

NR_046387.1_Zebra      aaaaaaaagcagcatggtgccaatcataaaaaaaaaa
NR_026757.1_Human      -------------------------------------
NR_015473.1_Mouse      -------------------------------------
```
