## Supplementary Materials for "LncRNA *OIP5-AS1* is overexpressed in undifferentiated oral tumors and integrated analysis identifies as a downstream effector of stemness-associated transcription factors"

### Table S12. List of Universal reverse transcription primers used for cDNA synthesis

| Study type | Primer type | Sequence |
|---|---|---|
| LncRNA | Universal RT primer | 5`CAGTGCAGGGTCCGAGGTACAGAGCCACCTGGGC AATTTTTTTTTTTVN-3` |
| | Universal 2nd strand reverse primer | 5`-CAGTGCAGGGTCCGAGGT-3` |
| miRNA | Stem loop RT primer – RNU44 specific | 5`-GTCGTATCCAGTGCGTCGAGTGACACGAGAGCCA CCTGGGCAATTTGCACTGGATACGACAGTCAG- 3` |
| | Stem loop RT primer – miR-30b-5p specific | 5`-GTCGTATCCAGTGCGTCGAGTGACACGAGAGCCA CCTGGGCAATTTGCACTGGATACGACAGCTGA- 3` |
| | Stem loop RT primer – miR-30a-5p specific | 5`-TCGTATCCAGTGCGTCGAGTGACACGAGAGCCAC CTGGGCAATTTGCACTGGATACGACCTTCCA- 3` |
| | Stem loop RT primer – miR-338-3p specific | 5`-TCGTATCCAGTGCGTCGAGTGACACGAGAGCCAC CTGGGCAATTTGCACTGGATACGACCAACAA- 3` |
| | Stem loop RT primer – miR-22-3p specific | 5`-TCGTATCCAGTGCGTCGAGTGACACGAGAGCCAC CTGGGCAATTTGCACTGGATACGACACAGTT- 3` |
| | Stem loop RT primer – miR-140-5p specific | 5`-TCGTATCCAGTGCGTCGAGTGACACGAGAGCCAC CTGGGCAATTTGCACTGGATACGACCTACCA- 3` |
| | Stem loop RT primer – miR-137 specific | 5`-TCGTATCCAGTGCGTCGAGTGACACGAGAGCCAC CTGGGCAATTTGCACTGGATACGACCTACGC- 3` |
| | Stem loop RT primer – miR-148a-3p specific | 5`-TCGTATCCAGTGCGTCGAGTGACACGAGAGCCAC CTGGGCAATTTGCACTGGATACGAC/ACAAAG- 3` |
| | Universal 2nd strand reverse primer | 5`-TCGTATCCAGTGCGTCGAGT-3` |
| mRNA | Random hexamer primers | 5`-NNNNNN-3` |

3` Wobble bases: V- [A,C,G], N- [A,C,G,T]

### Table S13. List of gene specific forward primers used for real time PCR experiments

| ncRNAs | Forward primer sequences |
|---|---|
| *GAPDH* | 5` - GAAGAGGGGAGGGGCCTAGG - 3` |
| *OIP5-AS1* | 5'- GCTTCCAAATCAGCAGAGGACCAT - 3' |
| *HOTAIR* | 5` - CTTGTGTAGGTTGTGTGTGTGTGGTGG - 3` |
| *NEAT1* | 5` - TCTTCTTCCCCTTTACAGCACAAAT - 3` |
| *TUG1* | 5' - GGCCGAGCGAACATGAAACTTTCAACT -3` |
| *RNU44* | 5`- GCAAATGCTGACTGAACATGA - 3` |
| miR-30b-5p | 5`- GCAGTGTAAACATCCTACACTCA - 3` |
| miR-30a-5p | 5`- GCAGTGTAAACATCCTCGACT - 3` |
| miR-338-3p | 5`- GCAGTCCAGCATCAGTGA - 3` |
| miR-22-3p | 5`- AGCTGCCAGTTGAAGAAC - 3` |
| miR-140-5p | 5`- CAGCAGTGGTTTTACCCTATG - 3` |
| miR-148a-3p | 5` - CAGTCAGTGCACTACAGAACT - 3` |
| miR-137 | 5`- CGCAGTTATTGCTTAAGAATACG - 3` |

**Table S14. List of primers used for SYBR® Green gene expression assays**

| Gene | Forward | Reverse |
|---|---|---|
| *GAPDH* | 5` - AGGGCTGCTTTTAACTCTGGT - 3` | 5` - CCCCACTTGATTTGGAGGGA - 3` |
| *CELF1* | 5` - CTGGACCACCCAGACCAACCA - 3` | 5` - CATGCATCCCTGGGAGGACTTTCA - 3` |
| *KMT2A* | 5` - CATCACCAGACCGACCTCCTCA - 3` | 5` - GGACCGCTGGGGTGATAAGGAA -3` |
| *KMT2C* | 5` - GGCTCATCACCGTTGTGTGGAGT - 3` | 5` - GGGCTGTCGCACACTGCAC - 3` |
