## Supplementary Materials for "LncRNA *OIP5-AS1* is overexpressed in undifferentiated oral tumors and integrated analysis identifies as a downstream effector of stemness-associated transcription factors"

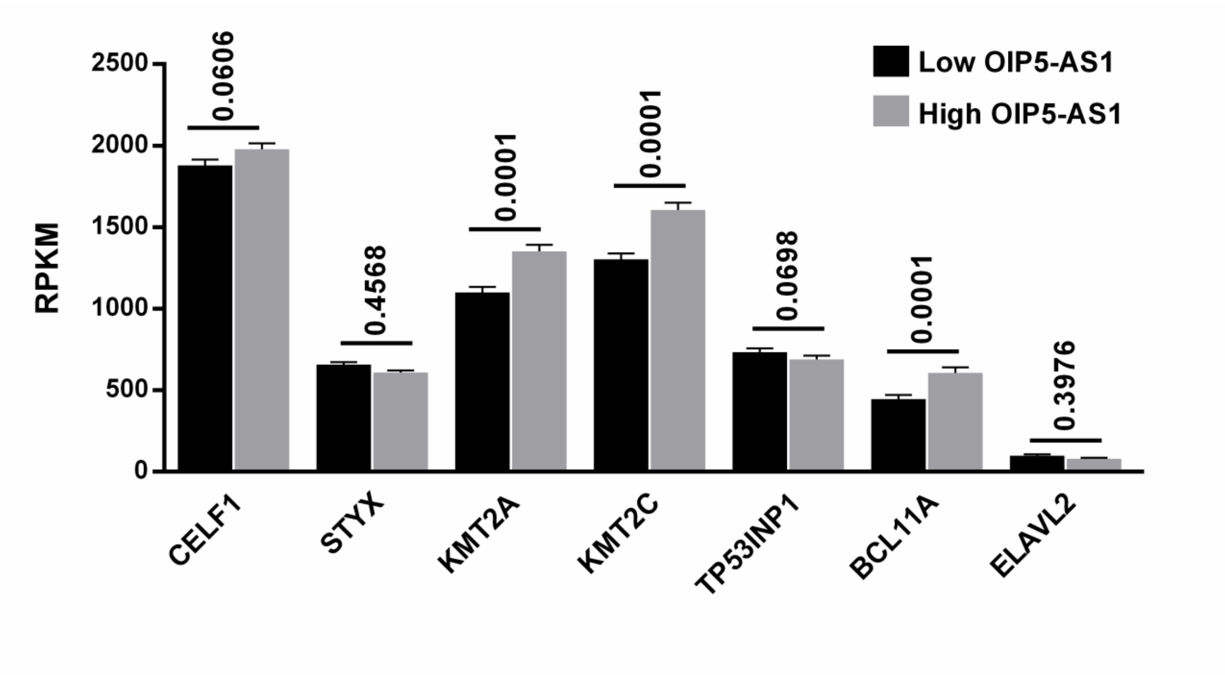


**Figure S2: Expression of candidate genes with lncRNA *OIP5-AS1* levels.** *KMT2A, KMT2C* and *BCL11A* were significantly upregulated in *OIP5-AS1* overexpressed HNSCC datasets from TCGA database. *CELF1* also showed overexpression with higher levels of *OIP5-AS1* in HNSCC.

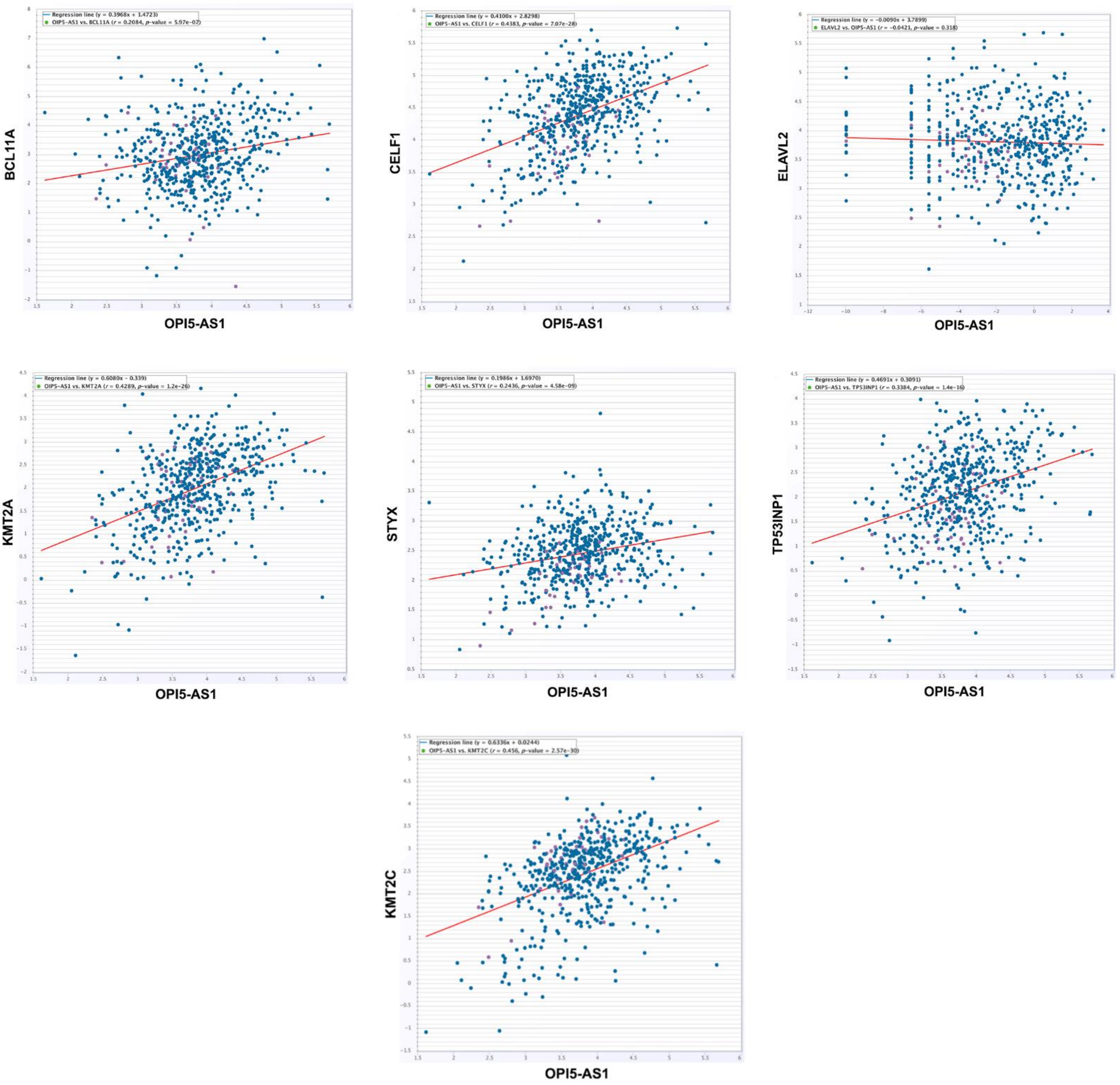


**Figure S3: Correlation of candidate genes with *OIP5-AS1* expression.** *CELF1*, *KMT2A* and *KMT2C* are having significant correlation with *OIP5-AS1* expression in head and neck cancer datasets from TCGA.

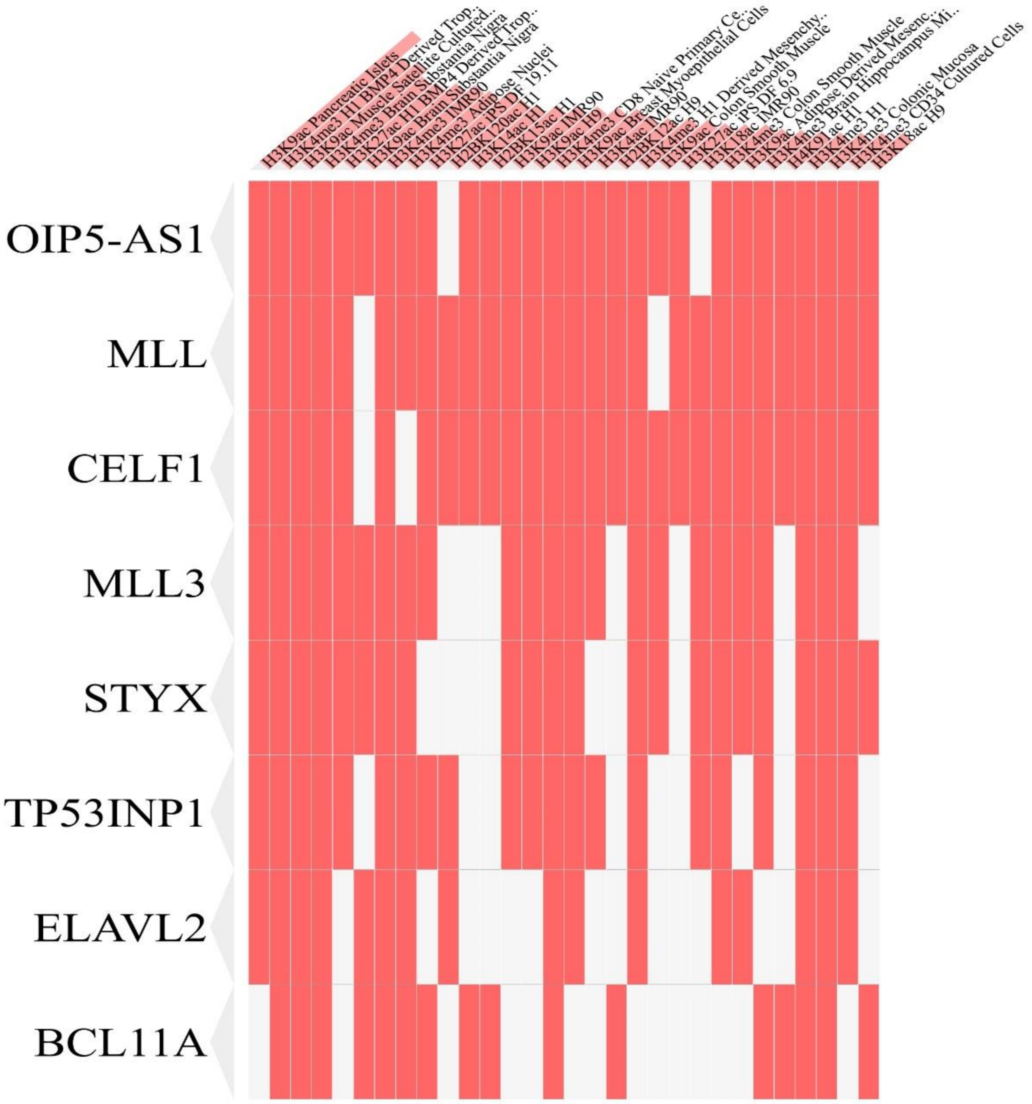


**Figure S4: Chromatin modifications at candidate gene locus in human cell types.** List of top 30 chromatin modifications at candidate gene locus in various human cell types by HM ChIP-seq from Roadmap Epigenomics Project. Along with lncRNA *OIP5-AS1* gene loci, *KMT2A, CELF1* and *KMT2C* are having significant active chromatin signature.

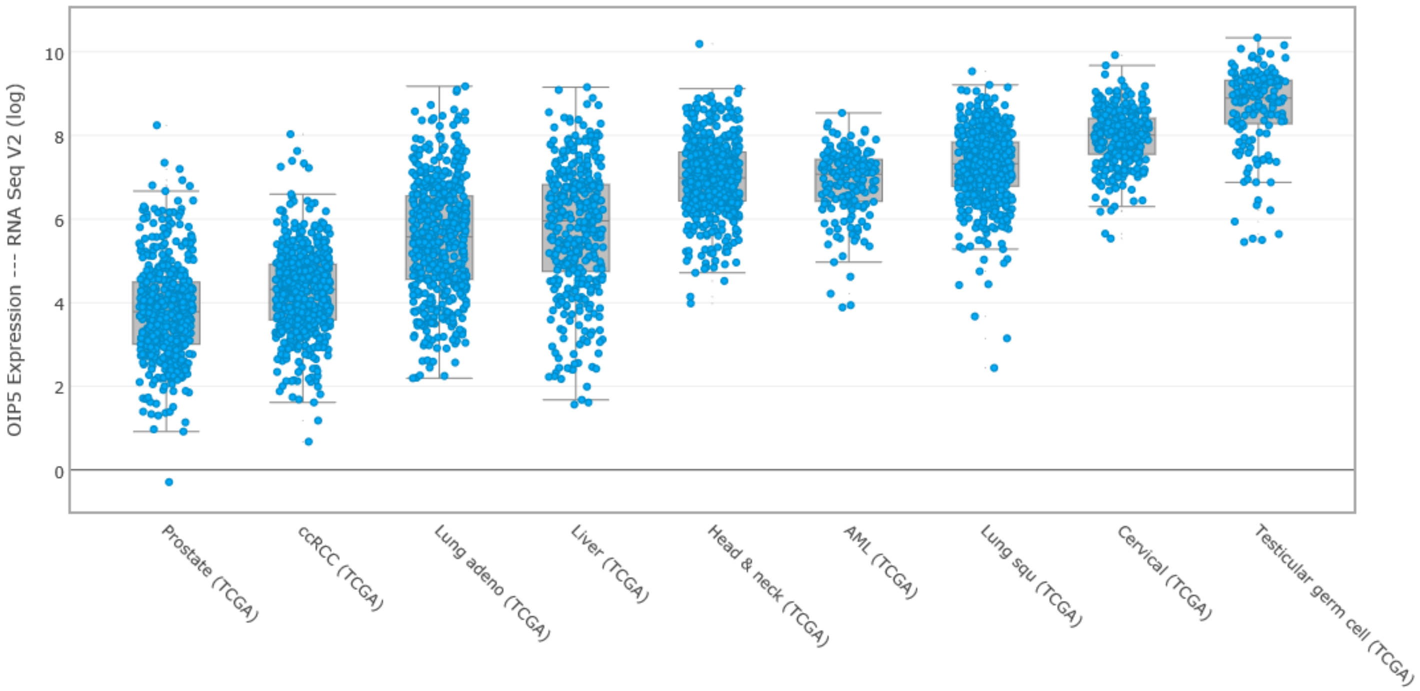


**Figure S5: Expression of *OIP5* across different type of cancers from TCGA dataset.** *OIP5* was significantly overexpressed in tumors of epithelial origin like *OIP5-AS1*. Testicular germ cell tumors expressed *OIP5* at very high level in comparison to all other tumors suggesting that *OIP5* also have a role in rapidly proliferating cells.

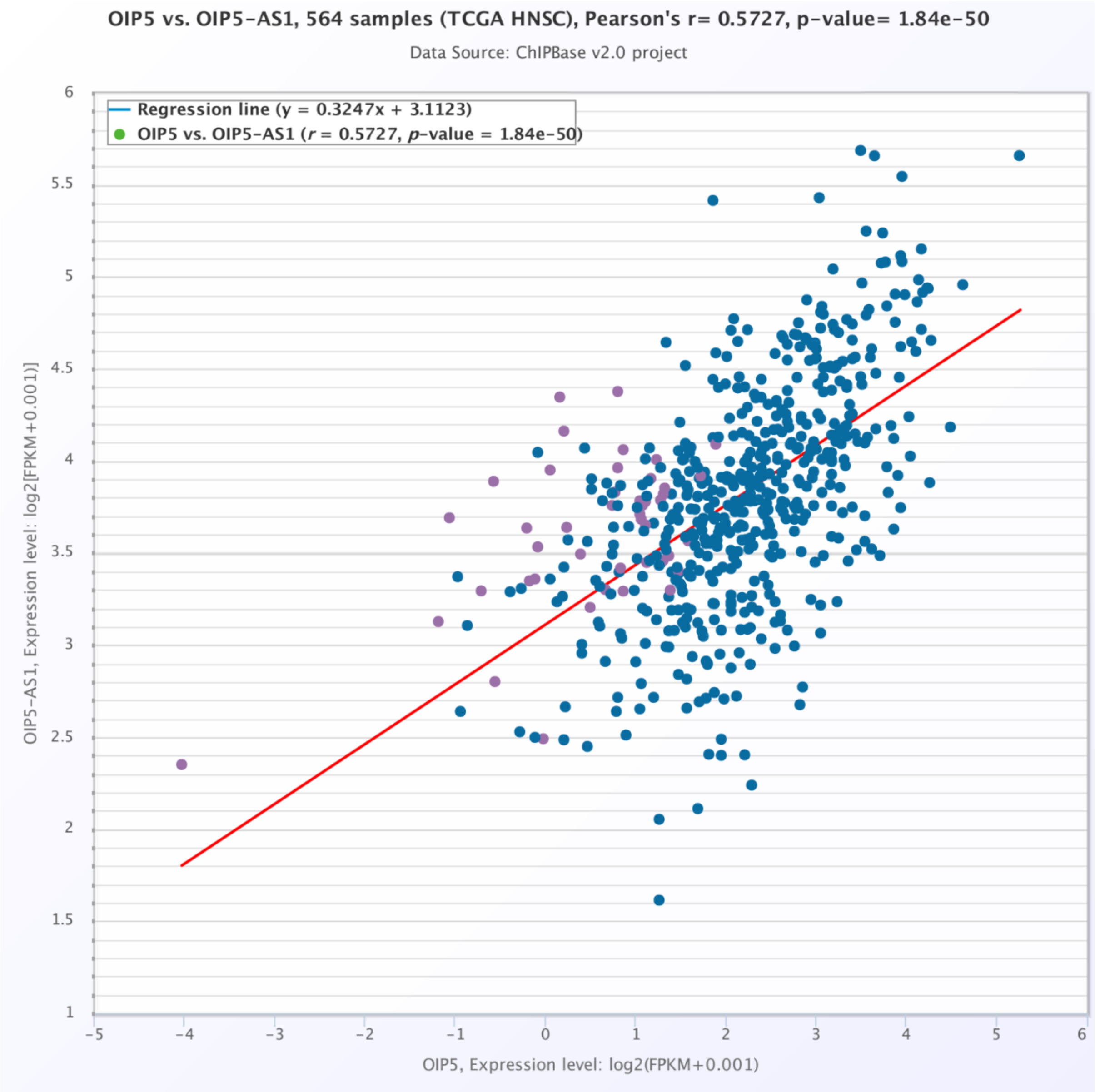


**Figure S6: Correlation plot of *OIP5* with lncRNA *OIP5-AS1* expression.** *OIP5* is the top significant co-expressed genes with LncRNA *OIP5-AS1* in HNSCC datasets of TCGA database ($r$ = 0.5727).
