## Supplementary Materials for "LncRNA *OIP5-AS1* is overexpressed in undifferentiated oral tumors and integrated analysis identifies as a downstream effector of stemness-associated transcription factors"

**Table S4: Functions of predicted miRNAs in various cancers.**

| Predicted miRNAs | Cancer types | Function/mechanism | Reference |
|---|---|---|---|
| hsa-miR-148a-3p | Gastric cancer | Tumor suppressor | PMID: 23456798 |
| | Colorectal cancer | Poor overall survival | PMID: 23933284 |
| | Pancreatic cancer | DNA hypermethylation | PMID: 20431052 |
| | Hepatocellular carcinoma | Inhibits metastasis | PMID: 24798342 |
| | Esophageal cancer | Recurrence/Survival | PMID: 20628822 |
| | Breast cancer | Tumor suppressor | PMID: 23554686 |
| | Ovarian cancer | Inhibits cell proliferation | PMID: 21971665 |
| | Head and neck cancers | Inhibits metastasis | PMID: 18768788 |
| hsa-miR-30a-5p | Lung adenocarcinoma | Inhibits tumor cell migration and invasion | PMID: 26837415 |
| | Non-small cell lung cancer | Sensitizes radio-therapy | PMID: 28259977 |
| | Ovarian cancer | Inhibits proliferation and invasion | PMID: 26675258 |
| | Hepatocellular carcinoma | Inhibits tumor proliferation and promotes apoptosis | PMID: 26884832 |
| | Head and neck cancers | Inhibits tumor cell progression | PMID: 26472042 |
| hsa-miR-30b-5p | Gastric cancer | Tumor suppressor | PMID: 25170877 |
| | Non-small cell lung cancer | Inhibits invasion and migration | PMID: 26388700 |
| | | Inhibits cell proliferation | PMID: 25249344 |
| | Laryngeal carcinoma | Promotes p53 induced apoptosis | PMID: 25356506 |
| | Breast cancer | Inhibits tumor cell growth and proliferation | PMID: 22384020 |
| hsa-miR-338-3p | Glioblastoma | Inhibits tumor proliferation | PMID: 28493990 |
| | Gastric cancer | Regulates EMT | PMID: 25945841 |
| | Non-small cell lung cancer | Proliferation and apoptosis | PMID: 28428733 |
| | Hepatocellular carcinoma | Inhibits cell growth and drug sensitization | PMID: 25531114 |
| | Ovarian cancer | Inhibits cell proliferation and metabolism | PMID: 27508048 |
| | Nasopharyngeal carcinoma | Inhibits tumor proliferation and migration | PMID: 26260688 |
| hsa-miR-22-3p | Hepatocellular carcinoma | Inhibits cell proliferation | PMID: 27904693 |
| | | Post-transcriptional regulation | PMID: 28045918 |
| hsa-miR-140-5p | Hepatocellular carcinoma | Cell growth and metastasis | PMID: 28383568 |
| | Osteosarcoma | Inhibits tumor proliferation and signals autophagy | PMID: 27582507 |
| | Cervical cancer | Growth arrest and inhibits metastasis | PMID: 27588393 |
| | Breast cancer | Prevents tumor invasion | PMID: 25983620 |
| | Hypopharyngeal carcinoma | Prevents tumor migration and invasion | PMID: 27033573 |
| hsa-miR-148b-3p | Non-small cell lung cancer | Tumor suppressor | PMID: 25232379 |
| | | Radio-sensitization | PMID: 26759383 |
| | Gastric cancer | Tumour cell metabolism | PMID: 28440026 |
| | Breast cancer | Context-dependent tumor suppressive | PMID: 25630670 |

| | | | |
|---|---|---|---|
| | | function | |
| hsa-miR-129-5p | Breast cancer | Regulates EMT | PMID: 26460733 |
| | Lung cancer | Inhibits cell proliferation and tumor invasion | PMID: 28105223 |
| | Gastric cancer | Reverses multi-drug resistance | PMID: 25344911 |
| | Laryngeal carcinoma | Growth arrest and apoptosis | PMID: 24194897 |
| | Hepatocellular carcinoma | Disease progression | PMID: 22536440 |
| | Ovarian cancer | Inhibits cell proliferation and survival | PMID: 25895125 |
| | Cervical cancer | anti-HPV activity | PMID: 24358111 |
| hsa-miR-137 | Neuroblastoma | Reverses drug resistance | PMID: 23934188 |
| | Colorectal cancer | Inhibits cell proliferation | PMID: 27764771<br>PMID: 23275153 |
| | Colorectal-Adeno carcinoma | Inhibits tumor progression | PMID: 28291253 |
| | Ovarian cancer | Inhibit EMT and invasion | PMID: 27596137 |
| | Breast cancer | Impairs tumor proliferative and migration | PMID: 22723937 |
| | Renal cell carcinoma | Tumor suppressor | PMID: 27347205 |
| | Lung cancer | Inhibits tumor growth and sensitizes chemosensitivity | PMID: 26989074 |
| hsa-miR-30e-5p | Colorectal cancer | Inhibits invasion and metastasis | PMID: 28656629 |
| | Breast cancer | Promotes proliferation, migration and invasion | PMID: 28288641 |
| hsa-miR-363-3p | Head and neck cancers | Reduces cell migration | PMID: 26545583 |
| | Lung adenocarcinoma | Inhibits tumor growth | PMID: 28423618 |
| | Gastric cancer | Inhibits cell growth and migration | PMID: 26709677 |
| | Papillary thyroid carcinoma | Inhibits tumor proliferation, migration and invasion | PMID: 28123856 |
| | Gallbladder cancer | Controls tumor progression | PMID: 27420766 |
| hsa-miR-424-5p | Neuroblastoma | Regulates ALK expression | PMID: 28455988 |
| | Cervical cancer | Inhibits tumor cell growth | PMID: 28082020 |
| | Gastric cancer | Promotes cell proliferation | PMID: 27655675 |
| | Esophageal cancer | Prevents tumor invasion and metastasis | PMID: 27628042 |
| | Non-small cell lung cancer | Inhibits proliferation, migration, and invasion | PMID: 27500472 |
| | Oral cancer | Induce cell migration and invasion | PMID: 27038552 |
| | Hepatocellular carcinoma | Inhibits cancer progression and EMT | PMID: 25175916 |
| | Pancreatic cancer | Promotes tumor proliferation, migration and invasion | PMID: 23653113 |
